# Balancing Stability and Flow in Hippocampal Networks via Inductive Bias and Learned Symmetry Breaking

**DOI:** 10.64898/2026.02.06.704443

**Authors:** Margot Wagner, Yusi Chen, Arjun Karuvally, Mia Cameron, Terrence J. Sejnowski

**Affiliations:** Institute for Neural Computation, University of California San Diego, La Jolla, CA; Institute of Neuroscience, Chinese Academy of Sciences, Shanghai, China; Computational Neurobiology Laboratory, Salk Institute for Biological Studies, La Jolla, CA; Computational Neuroscience Center, University of Washington, Seattle, WA

**Keywords:** hippocampus, RNN, CA3, sequence generation, replay, prediction, symmetry breaking, inductive bias

## Abstract

The hippocampus must balance stable memory representations with internally generated sequential dynamics underlying replay and prediction. How hippocampal circuitry achieves both remains unclear. Here, we show that recurrent neural networks trained on prediction tasks converge to a mixed-symmetry dynamical regime, in which dominant symmetric recurrence stabilizes an attractor while a weaker antisymmetric component induces directed flow. This structure accelerates learning and supports robust replay and prediction. We further show that such symmetry breaking can arise from biologically plausible spike-timing-dependent plasticity (STDP) rules, yielding a tilted Mexican-hat connectivity profile without explicit architectural constraints. Initializing networks with CA3-like structured connectivity biases learning toward this regime, and improves optimization efficiency and performance. These results suggest that hippocampal computation reflects an interaction between biologically constrained circuit structure and task-driven learning, with partial symmetry breaking providing a low-dimensional control principle for balancing stability and flow in sequence generation.

**Subject Areas:** computational neuroscience, systems neuroscience, neural dynamics, learning, memory

## 1. Introduction

### (a) Hippocampal memory: stable attractors and directed sequential activation

The hippocampus plays a central role in learning, storing, and retrieving sequential experience. From early spatial map theories [1] to contemporary views of the hippocampus as a generator of predictive internal models [2,3], a central challenge has been to reconcile memory stability with temporally ordered neural activity. In particular, how can neural circuits preserve memories while also supporting structured sequences over time? The hippocampus and especially its sub-region CA3 have long been proposed as recurrent associative memory systems [4–6] capable of rapid pattern completion through plastic recurrence. In these theories, CA3 operates as an autoassociative attractor network in which recurrent excitation stabilizes stored representations and enables pattern completion from partial cues [7–10]. While this model explains stable memory retrieval, accumulating experimental and theoretical evidence suggests that hippocampal activity also exhibits intrinsically sequential dynamics [11–16]. Rather than simply reinstating stored representations, hippocampal activity frequently unfolds as internally generated sequences, implying intrinsic temporal organization beyond static recall [17–23]. These observations raise a central question of how hippocampal circuits can maintain stable attractor-like representations while also supporting directed sequential activity.

Converging experimental evidence suggests that hippocampal population dynamics are also often inherently sequential and directional. During both sleep and awake behavior, hippocampal populations exhibit temporally ordered reactivation of place cell sequences of behaviorally relevant past and future trajectories [6,14,17,24–30]. These replay events reflect structured progression through internal representations rather than noise-driven fluctuations around a fixed point and are thought to support prediction and planning [14,15,18]. Theta phase precession, traveling waves, and directionally biased sequences further indicate that hippocampal circuits support both representational persistence and controlled temporal progression [11–13,19,22,31]. These observations motivate a closer examination of classical hippocampal recurrent network models.

In addressing this question, here the focus is on population-level modeling. While continuous-rate recurrent neural networks (RNNs) provide a tractable framework for studying low-dimensional attractor structure, they abstract away the sparse firing of hippocampal pyramidal cells. The relationship between firing-rate attractor models and low-rate spiking networks remains an open theoretical question [32–34]. However, prior work and related simulations using the population activity show that spiking networks can reproduce key qualitative dynamics of trained rate networks, including replay and phase precession, under appropriate conditions [2,35]. The present work focuses on dynamics at the population level rather than detailed spiking networks.

Classical theories of recurrent neural computation emphasize symmetry as a prerequisite for stable memory. Fully symmetric recurrent networks converge to fixed-point attractors, providing a principled mechanism for content-addressable associative memory and pattern completion [32,33,36,37]. This idea strongly influenced computational theories of hippocampal area CA3, which has long been modeled as a recurrent, symmetric autoassociative network optimized for rapid storage and retrieval of episodic patterns [4,33]. Related theoretical work has examined how specific recurrent connectivity motifs support attractor memory storage and recall, predicting structured patterns of reciprocal connectivity in networks optimized for associative memory [38]. Extensions of this approach to continuous attractor networks, including ring and bump models, demonstrated how symmetric recurrent excitation combined with broader inhibition (Mexican-hat) can stabilize low-dimensional manifolds representing continuous variables, yielding explanations for spatial coding, path integration, and head-direction representations [21,34,39–42]. Despite their success in explaining persistent activity and pattern completion, classical symmetric attractor models do not by themselves generate directed sequential dynamics. Introducing asymmetric recurrent interactions enables temporal transitions between memory states [14,39,43,44].

Early models of sequence generation in recurrent networks proposed that stable attractor dynamics can coexist with slower transition dynamics that move activity between states. For example, Sompolinsky and Kanter [44] and Kleinfeld [43] showed that combining symmetric memory couplings with asymmetric interactions or delayed synaptic response produces sequences of long-lived quasi-equilibrium states, while Lisman [14] extended this framework to hippocampal circuits by proposing a separation between autoassociative stabilization and heteroassociative progression. Related models suggested that sequences are explicitly stored through temporally asymmetric plasticity using transitions between discrete activity patterns [13,45]. Other work emphasized oscillatory mechanisms in which theta rhythms organize sequential firing and phase precession via phase coding or theta-gamma segmentation [46–49].

In addition to these mechanisms, several models have proposed that sequential dynamics arise from velocity-driven transitions along low-dimensional manifolds, in which external motion-related signals drive activity through representational space [50–52]. These approaches provide a compelling account of sequence generation during navigation and path integration. However, they rely on externally supplied velocity signals to drive transitions between states, leaving open the question of how sequential structure may be generated intrinsically within recurrent circuitry, particularly during offline states such as replay.

In contrast to these approaches which rely on transitions between discrete attractor states, externally imposed temporal structure, or transient dynamical mechanisms such as adaptation or oscillatory coupling, the present model shows that sequential dynamics can emerge intrinsically from learned recurrent connectivity. Specifically, training drives the network toward a regime in which dominant symmetric structure stabilizes a continuous attractor manifold, while a weaker, geometrically aligned antisymmetric component induces directed flow along that manifold. In this regime, replay and prediction arise as smooth, internally generated trajectories rather than discrete state transitions, without requiring explicitly stored transitions, oscillatory discretization, or hand-crafted dynamical asymmetries.

Heteroassociative chains can support sequential recall by linking successive states through asymmetric connections. However, classical analyses of CA3 networks emphasize the role of autoassociative connectivity in enabling pattern completion and stabilizing stored representations in the presence of noise or partial cues [4,14]. In these models, symmetric recurrent interactions allow the network to recover a stored representation before sequential transitions occur. Purely heteroassociative chains lack this stabilizing mechanism and can therefore be sensitive to noise and error accumulation. The regime explored here can be interpreted as a hybrid of these mechanisms, allowing stable representations and sequential dynamics to coexist within a single recurrent circuit [14,43,44]. In these models, asymmetry plays a functional role in generating sequences. However, when not structured relative to the underlying attractor geometry, increasing asymmetry can reduce the stability or persistence of individual states, leading to a tradeoff between memory stability and directed transitions. This highlights a fundamental balance: while symmetry supports memory stability, directionality is required for sequence generation, leaving open the question of how hippocampal circuits balance these competing demands. This suggests that hippocampal circuits may operate in an intermediate dynamical regime that balances stability and directed flow.

### (b) The need to naturally break symmetry through neural mechanisms

Controlled symmetry breaking may link stable memory and directed temporal dynamics. In continuous attractor models, symmetric (even) recurrent interactions stabilize a low-dimensional representational manifold, while a carefully structured antisymmetric (odd) component induces smooth motion along that manifold without distorting the underlying representational structure [53]. Even modest asymmetry can convert a static attractor into a system capable of controlled flow. This approach has been used to explain head-direction integration, phase precession, and traveling population activity, in which weak antisymmetric or non-reciprocal interactions generate drift or rotation atop a stabilizing recurrent backbone [13,19,20,53–56]. Crucially, the structure of asymmetry matters: antisymmetric components aligned with spatial derivatives preserve representational structure and produce coherent translation, whereas unstructured asymmetry degrades stability or induces chaotic dynamics [44,57].

In hippocampal models, directed dynamics have been attributed to several sources of symmetry breaking, including anatomically biased recurrent connectivity, temporally asymmetric plasticity, and transient dynamical effects such as short-term synaptic processes [14,21,54,58–65]. Experimental observations are consistent this view, showing traveling hippocampal activity consistent with internally generated flow along representational axes [19,56,66]. These results highlight a central problem: hippocampal circuits must break symmetry to support replay and prediction without sacrificing stability.

One biologically plausible route to controlled symmetry breaking is synaptic plasticity. In particular, mixed symmetry in recurrent CA3 connectivity may emerge from diverse spike-timing-dependent plasticity (STDP) rules across hippocampal synapses. Early formulations of STDP emphasize a temporally asymmetric learning window in which pre-before-post activity potentiates synapses and post-before-pre activity depresses them, establishing STDP as a causality-sensitive Hebbian mechanism [67,68]. However, the CA3 may not be governed solely by canonical STDP rules.

Direct measurements at CA3-CA3 recurrent excitatory synapses reveal a largely symmetric STDP rule, well matched to CA3 autoassociative memory storage and recall [69]. Theoretical work also suggests that temporally asymmetric plasticity can coexist with more symmetric components in hippocampal-like circuits, jointly supporting pattern completion and temporal prediction [58]. Recent theory further formalizes this heterogeneity, showing that subpopulations with differing degrees of temporal asymmetry in their learning rules can regulate replay speed and directionality, with symmetric components stabilizing activity and asymmetric components driving transitions [70]. Emerging CA3 microcircuit data also indicate structural and subtype-specific asymmetries in recurrent excitation that may interact with heterogeneous plasticity to support both stable attractor dynamics and controlled sequence generation [69,71–73].

### (c) Inductive biases in hippocampal RNN models

The hippocampus is frequently modeled as a recurrent neural network (RNN) due to its recurrence, plasticity, and central role in sequence learning and prediction [2,58,59,74]. A central unresolved question is whether hippocampal-like dynamics arise primarily through learning from initially unstructured connectivity, or whether circuit-level inductive biases imposed by development or evolution constrain the space of attainable dynamics. Recent RNN studies have demonstrated that predictive and sequential structure can, in principle, emerge from training networks initialized with random recurrent weights [2,54,57,74,75]. Yet because random connectivity lacks biological specificity, it remains unclear which learned features are essential for computation and which are incidental. In contrast, converging experimental and theoretical work suggests that hippocampal area CA3 is not randomly wired, but instead exhibits structured, distance-dependent recurrent connectivity that supports coordinated neural assemblies and continuous attractor dynamics [21,37,76,77]. Canonical center-surround (Mexican-hat-like) connectivity stabilizes representations through symmetric interactions, while modest symmetry breaking has long been predicted to enable directed flow [14,20,43,44,53]. These considerations motivate a systematic examination of whether biologically motivated recurrent structure shapes learning efficiency, dynamical stability, and the emergence of replay and prediction.

### (d) Contribution

A central open question in hippocampal theory is how recurrent circuits stabilize internal representations while generating ordered temporal dynamics. We hypothesize that hippocampal CA3 operates in an intermediate regime where symmetric connectivity stabilizes representations while a weaker antisymmetric component drives sequential propagation. Here, we examine how the balance between symmetric and antisymmetric recurrent connectivity governs learning, replay, and prediction in CA3-like recurrent networks.

First, using an Elman-style RNN trained on a circular prediction task inspired by place-cell navigation, we show that networks trained from random connectivity reliably self-organize into a structured regime dominated by a symmetric Mexican-hat-like component with a smaller but functionally essential antisymmetric contribution, which we term a “tilted Mexican hat.” This mixed-symmetry organization emerges through learning and supports stable attractors while generating sequential dynamics underlying replay and prediction (Figure 1).

**Figure 1:**
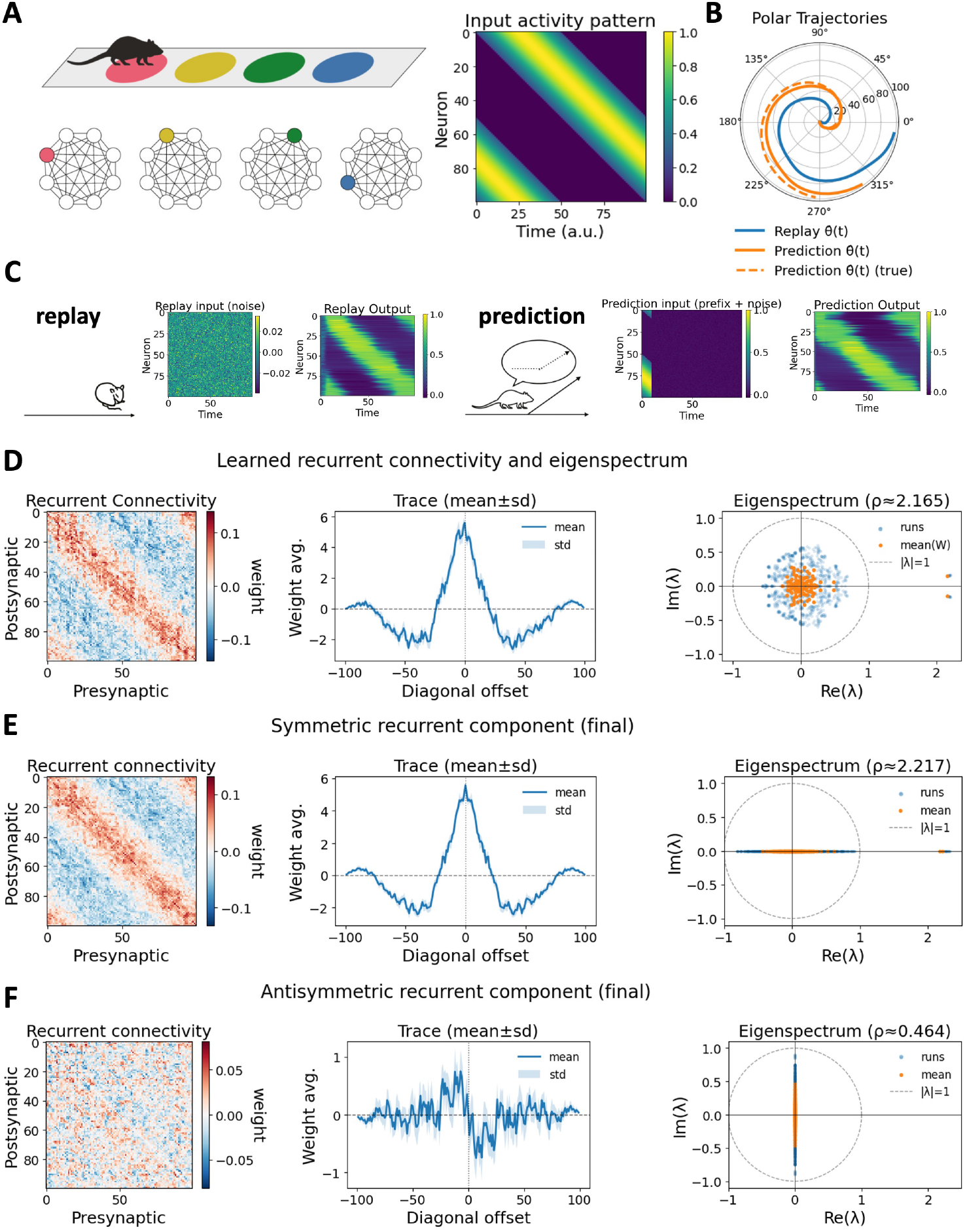
Learned symmetry breaking in recurrent connectivity supports replay and prediction. **(A)** Task schematic: a rat traverses a circular track and receives bell-shaped, location-specific input. **(B)** Polar trajectories during replay (*blue*) and prediction (*orange*), showing coherent sequential dynamics. **(C)** Evaluation paradigms: replay driven by noise (*left*) and prediction following a brief input prefix (*right*). **(D)** Learned recurrent weight matrix, averaged across runs and sorted by neurons’ peak activation time, revealing a banded structure aligned with sequence order. **(E)** Symmetric component of the learned connectivity, exhibiting a Mexican-hat-like profile and real-valued eigenmodes. **(F)** Antisymmetric component, showing off-diagonal structure and imaginary eigenmodes consistent with directional or flow-like dynamics. Eigenspectra in panels D-F are computed from the recurrent weight matrix *W*_*hh*_ and are interpreted as structural diagnostics of learned connectivity rather than direct stability criteria for the nonlinear dynamics.

Second, we show that biologically inspired CA3-like inductive biases improve both optimization and computation (Table 1 and 2). Tilted Mexican hat initialization accelerates early learning, reinstates a two-phase optimization regime absent under random connectivity, yields tighter alignment between replay and prediction dynamics, and uncovers a spatial derivative drift component. These results show that structured recurrent connectivity does not merely emerge post hoc, but constrains learning toward functionally relevant solutions.

**Table 1:** Replay metrics.

| metric | random | identity | pure shift | mixed shift | Mexican hat ($\alpha_0 = 0.7$) |
| --- | --- | --- | --- | --- | --- |
| ring decode $R^2$ (↑) | $0.963 \pm 0.012$ | $0.211 \pm 0.27$ | $0.782 \pm 0.013$ | $0.929 \pm 0.027$ | **$0.975 \pm 0.0029$** |
| residual lag-1 autocorr (↑) | $0.99 \pm 0.0032$ | $0.176 \pm 0.27$ | $0.99 \pm 0.0029$ | $0.99 \pm 0.002$ | **$0.99 \pm 0.0061$** |

**Table 2:** Prediction metrics.

| metric | random | identity | pure shift | mixed shift | Mexican hat ($\alpha_0 = 0.7$) |
| --- | --- | --- | --- | --- | --- |
| mean resultant length $R$ (↑) | $0.956 \pm 0.0071$ | $0.939 \pm 0.015$ | $0.836 \pm 0.0083$ | $0.922 \pm 0.024$ | **$0.987 \pm 0.0044$** |
| ring decode $R^2$ (↑) | $0.968 \pm 0.011$ | $0.957 \pm 0.014$ | $0.778 \pm 0.012$ | $0.921 \pm 0.036$ | **$0.99 \pm 0.0039$** |
| MSE (↓) | $0.0646 \pm 0.0066$ | $0.079 \pm 0.0076$ | $0.114 \pm 0.0052$ | $0.0814 \pm 0.013$ | **$0.0499 \pm 0.0046$** |

Third, we demonstrate that the balance between symmetric and antisymmetric connectivity constitutes a low-dimensional control parameter governing recurrent network dynamics. By explicitly manipulating the symmetry-antisymmetry ratio of structured recurrent weight matrices, we reveal transitions between static attractors under pure symmetry, unstable drift or oscillation under pure antisymmetry, and robust flow-stabilized attractors at intermediate values (Figure 2). Networks initialized near this intermediate regime learn faster, exhibit more reliable autonomous replay, and produce more accurate predictions than networks initialize with random, purely symmetric, or purely directed connectivity motifs.

**Figure 2:**
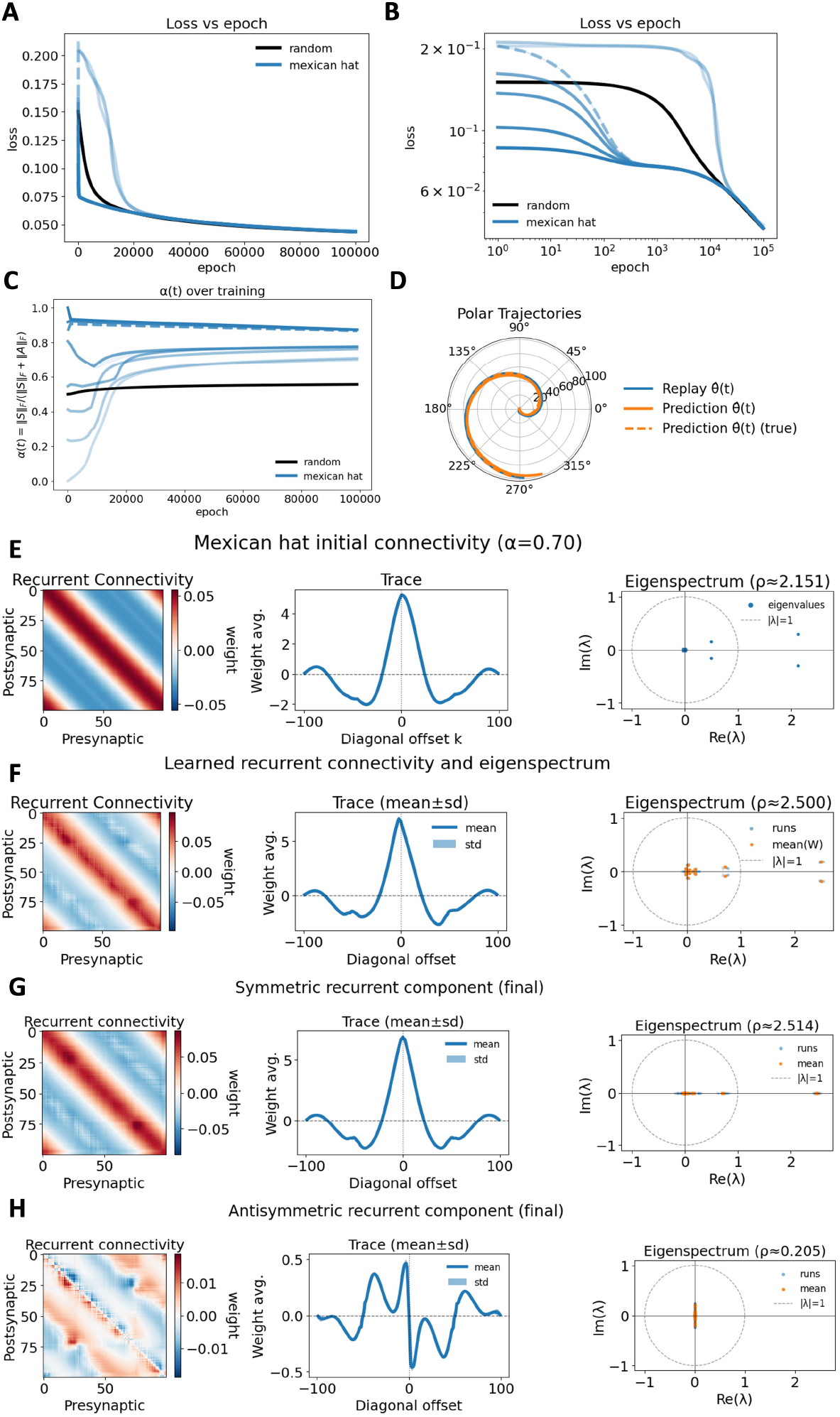
Tilted Mexican-hat CA3-like recurrent connectivity accelerates learning and stabilizes sequential dynamics. **(A**,**B)** Training loss for networks initialized with circulant Mexican-hat connectivity (blue; varying initial symmetry *α*_0_, dashed: *α*_0_ = 0.70) compared with random connectivity (black), shown on linear and log-log axes. **(C)** Evolution of the symmetry index *α*(*t*), showing convergence to an intermediate, symmetry-dominated regime. Initial symmetry *α*_0_ is given by value at epoch 0. **(D)** Polar trajectories of replay and one-step-ahead prediction for *α*_0_ = 0.70. **(E)** Initial Mexican-hat connectivity (*α*_0_ = 0.70), its diagonal trace, and eigenspectrum. **(F)** Learned recurrent connectivity after training. **(G**,**H)** Symmetric (G) and antisymmetric (H) components of the learned connectivity, showing a dominant attractor-like scaffold with a weaker directional component. Eigenspectra shown in panels E-H correspond to *W*_*hh*_. The effective Jacobian is contracted by the state-dependent tanh gain during replay and prediction.

Finally, we provide a biologically plausible synaptic mechanism for the emergence of mixed-symmetry connectivity. We show that a modified STDP rule combining predominantly symmetric plasticity with a weak antisymmetric component naturally produces tilted Mexican-hat connectivity profiles. This learning rule generates stable representational wave dynamics whose speed and direction depend continuously on the degree of symmetry breaking (Figure 3), linking classical STDP, recent experimental evidence for symmetric CA3-CA3 plasticity, and the emergence of directed hippocampal sequences.

**Figure 3:**
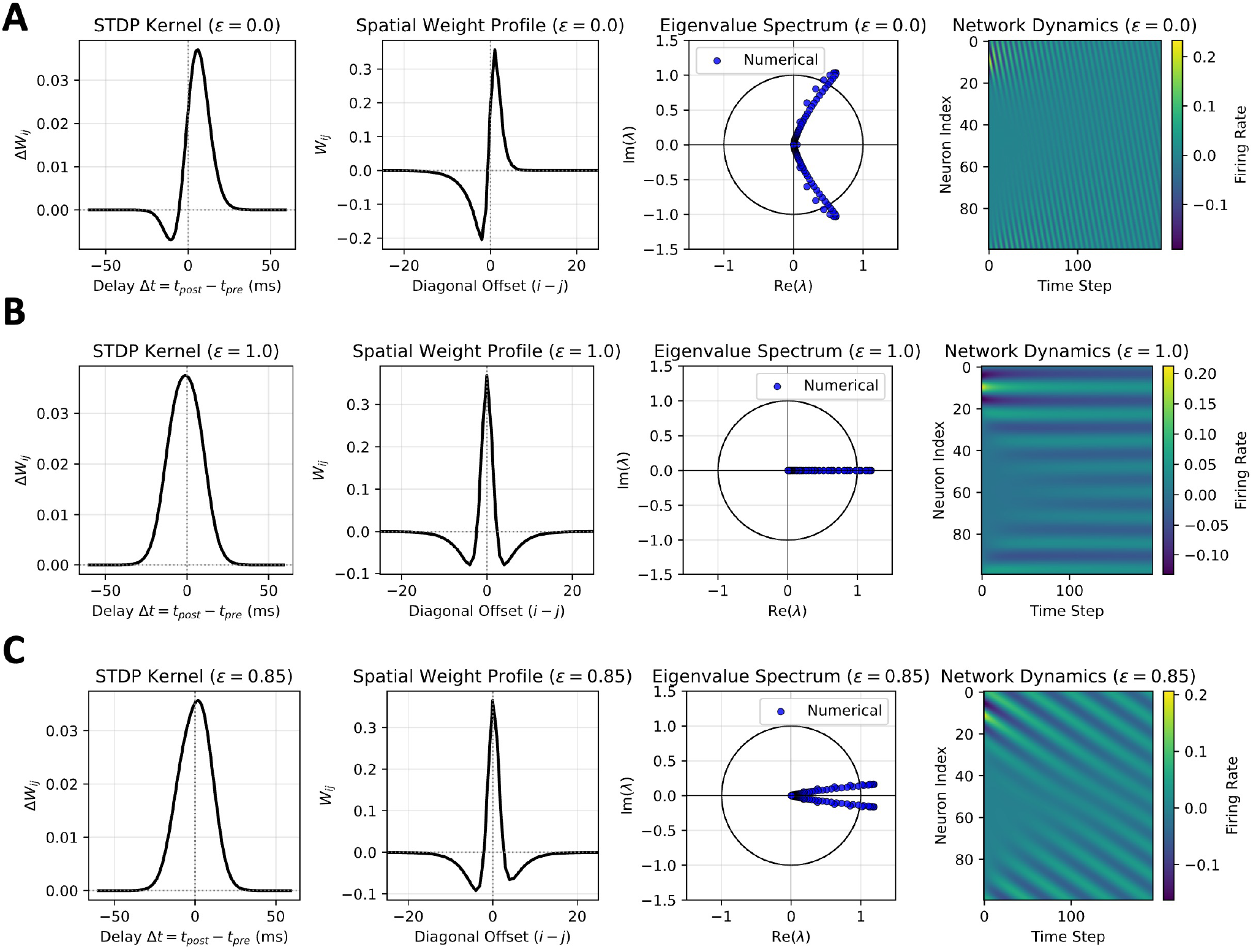
Symmetry breaking in STDP generates flow-stabilized Mexican-hat dynamics. Each panel shows the learned synaptic weight profile for different settings of the symmetry parameter (*ϵ*). The first column shows the STDP learning kernel, the second column the weight profile after learning, the third column the computed eigenvalue spectrum of the learned matrix and the fourth column the resulting network activity dynamics. We use STDP with parameters *A*_+_ = 1.0, *A*_−_ = 0.5, *τ*_+_ = 1.0, *τ*_−_ = 3.0. **(A) Pure symmetry** (*ϵ* = 1): symmetric Mexican-hat profile with real eigenvalues and a stationary attractor. Activity bump is stationary (zero velocity). **(B) Pure antisymmetry** (*ϵ* = 0): standard STDP gives highly asymmetric weight profile with fast activity drift. **(C) Symmetry breaking** (*ϵ* = 0.85): weakly tilted Mexican hat with mixed real-imaginary eigenvalues, producing a flow-stabilized attractor with a slowly propagating activity bump.

Our results support a view of the hippocampus as a flow-stabilized recurrent system operating in an intermediate dynamic regime, where symmetric recurrence stabilizes internal representations and structured antisymmetry generates controlled temporal progression. *More broadly, our findings suggest that hippocampal computation reflects an interaction between task-driven learning and circuit-level inductive biases, with symmetry breaking organizing replay and prediction*.

## 2. Methods

### (a) Task and network architecture

To examine the impact of structural inductive bias in hippocampal recurrent circuitry, we developed a predictive RNN model inspired by place-cell navigation. The task consisted of a simulated rat traversing a circular track at constant velocity. Neurons in the recurrent layer, representing CA3, received location-specific bell-shaped inputs that serve as a simplified encoding of spatial position, analogous to place fields, rather than a direct model of sensory-driven place-cell response (Fig. 1A). The network was trained to perform one-step-ahead prediction [2], mapping population activity at time *t* to activity at time *t* + 1. The recurrent units implement population-level firing-rate dynamics and do not explicitly model spike timing, synaptic conductances, or oscillatory interactions present in biological CA3 circuits. Throughout, the recurrent weight matrix is interpreted as an effective coupling in a reduced rate model. Because Dale’s principle is not enforced, negative weights represent effective net inhibitory influence at the population level.

The model was implemented as an Elman RNN in PyTorch (v2.6.0) and trained using gradient descent with backpropagation through time and a fixed learning rate. Recurrent units used a tanh nonlinearity, chosen for its bounded output and smooth derivative which facilitate stable training and recurrent dynamical analysis, and output units used a sigmoid nonlinearity. Input-to-hidden and hidden-to-output weights were frozen to isolate the role of recurrent dynamics. Training terminated when either a maximum epoch count was reached or the loss dropped below 1% of its initial value and changed by less than 10^−3^% over ten consecutive epochs. The network was first trained starting from a random initialization then using crafted structural hidden weights. The use of inductive bias for improved training was motivated by pretraining (see Fig. S1), but the hidden weights were initialized from different connectivity families. Network dimensions and hyperparameters are summarized in Table S1. Training sequences consisted of repeated laps around the circular track with fixed direction and velocity.

### (b) Recurrent connectivity families and inductive biases

We examined multiple recurrent connectivity families encoding distinct inductive biases: random, identity, cyclic shift, and Mexican-hat connectivity. Random initialization served as the baseline with other families normalized to match.

We explicitly define the connectivity families as follows. Let *W* ∈ ℝ denote the recurrent weight matrix. Identity connectivity is defined as

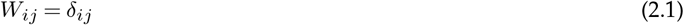

where *δ*_*ij*_ is the Kronecker delta. This corresponds to purely self-recurrent dynamics with no interactions, thereby isolating temporal persistence.

Cyclic shift connectivity is defined as

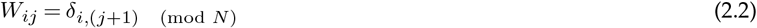

which implements a discrete translation operator on the ring, such that activity at neuron *j* is passed to neuron *j* + 1 with wrap-around. Although visually similar to identity (a diagonal matrix), cyclic shift connectivity is strictly off-diagonal, implementing directional propagation rather than persistence.

Both the cyclic shift and Mexican-hat matrices are circulant, meaning their rows are cyclic permutations of a base kernel, ensuring translation-invariant connectivity on the ring.

Further discussion and visualizations of all connectivity families (random, identity, cyclic shift, and Mexican-hat) are provided in the Supplemental Material 6(c) (see Fig. S2).

### (c) Symmetry-antisymmetry decomposition and control parameter

In addition to the baseline cyclic shift and Mexican-hat initializations, we also implemented variants with predetermined symmetry components. Specifically, the recurrent matrices *W* were decomposed into symmetric *S* and antisymmetric *A* components

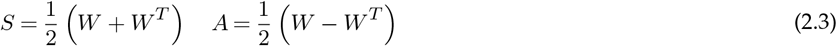

and recombined as

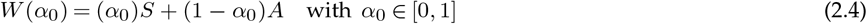

The parameter *α*_0_ controls the initial balance between reciprocal (symmetric, *α*_0_ = 1) and directional (antisymmetric, *α*_0_ = 0) interactions. To track how symmetry evolved during training, we defined a time-dependent symmetry index using Frobenius norms as

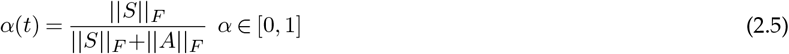

which provides a low-dimensional summary of recurrent structure, explicitly parameterizing the degree of symmetry breaking in the network. *α* increases as the symmetric component becomes dominant, where *α* = 0 represents the purely anti-symmetric extreme and *α* = 1 denotes the purely symmetric state.

### (d) Training, replay, and prediction protocols

Networks were trained using teacher forcing [78], with the true input provided at each time step. We used two evaluation regimes: prediction and replay. For replay, networks were driven entirely by low-amplitude noise to assess internally generated dynamics. For prediction, networks received a short prefix of true input followed by noise and were assessed on their ability to autonomously continue the learned sequence.

### (e) Mexican-hat connectivity construction

Mexican-hat connectivity was constructed by defining a one-dimensional Difference-of-Gaussians (DoG) kernel on a ring of size *N*. For offsets from the diagonal *d* ∈ 0, 1, …, *H* − 1, we define the signed circular distance as

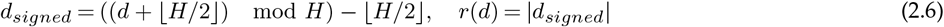

and construct the DoG kernel

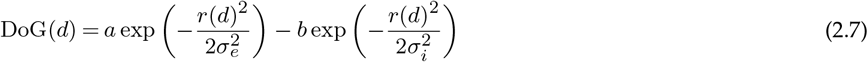

where *σ*_*e*_ and *σ*_*i*_ are the excitation and inhibition widths, and *a* and *b* are the excitatory and inhibitory amplitude respectively. *b* was chosen to approximately balance total excitation and inhibition. The DoG inhibitory lobe denotes an effective negative interaction in the rate model. Enforcing Dale’s principle would require an explicit excitatory-inhibitory architecture or sign-constrained parameterization, which is not imposed here. In rate-based network models, the recurrent weight matrix should be interpreted as an effective interaction operator rather than a direct representation of individual synaptic connections. The kernel was scaled to match a target spectral radius *ρ*_*target*_ using the maximum magnitude of its Fourier coefficients. In our experiments, *σ*_*i*_ *> σ*_*e*_ to produce a local excitatory bump with broader surround inhibition.

The final recurrent weight matrix was obtained by embedding the kernel into a circulant matrix

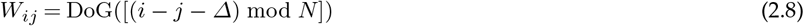

where Δ optionally introduces a directional shift. For shifted Mexican-hat matrices, the symmetry-antisymmetry balance was further controlled via the *α* decomposition described above.

### (f) Plasticity analysis

To identify a local synaptic mechanism capable of producing the mixed-symmetry connectivity for hippocampal sequence generation, we developed a modified spike-timing-dependent plasticity (STDP) rule [67]. This STDP rule completely replaces backpropagation in the final experiment. While canonical STDP is temporally asymmetric, strengthening synapses only when presynaptic activity precedes postsynaptic firing [68],this causal rule contradicts the dual role of the hippocampus in forming stable memory representations and enabling directed sequential dynamics.

Our rule is motivated by recent empirical evidence suggesting that CA3 recurrent circuitry is not governed solely by canonical, asymmetric STDP [69]. Direct electrophysiological measurements at CA3-CA3 recurrent excitatory synapses have revealed a predominantly symmetric STDP rule, where potentiation occurs for both pre-before-post and post-before-pre spike timings. This symmetric plasticity is thought to be functionally optimized for the rapid storage and stable retrieval of autoassociative memories. We therefore implement a rule that allows for the strengthening of “reverse” synapses (post-before-pre) during forward spiking events, effectively modeling the retroactive potentiation observed in biological CA3 ensembles.

We formalize this by modifying the canonical STDP such that it linearly interpolates between a standard antisymmetric STDP kernel, *K*(*s*), and a retroactive, symmetry-inducing component controlled by a scalar parameter *ϵ* ∈ [0, 1]. The total synaptic update from neuron *j* to neuron *i* is calculated by integrating the product of neural activities *u*(*t*) across a biphasic kernel:

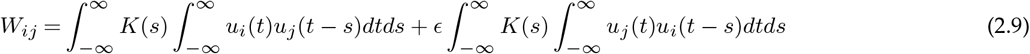

where the biphasic STDP kernel *K*(*s*) is defined as:

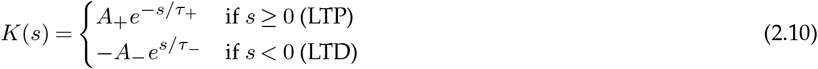

In these expressions, *A*_+_ and *A*_−_ represent the maximum learning rates, while *τ*_+_ and *τ*_−_ denote the time constants for potentiation and depression, respectively. The parameter *ϵ* serves as a control for the degree of symmetry breaking at the synaptic level. This formulation allows the circuit to interpolate between purely antisymmetric (*ϵ* = 0) and purely symmetric (*ϵ* = 1) regimes, providing an analytically tractable mechanism for the hippocampus to balance representational stability with controlled temporal progression through local changes in the retrograde plasticity signal. We then analyzed the emergent network structure when a simple linear RNN is stimulated with the moving Gaussian input activity profile of the form used in our experiments.

## 3. Results

### (a) Task-driven symmetry breaking in recurrent connectivity supports replay and prediction

Training an Elman recurrent neural network from random initialization was sufficient to generate hippocampal-like replay and one-step-ahead prediction on a circular navigation task (Fig. 1A-C). This is consistent with prior work showing that generic recurrent networks can support sequential dynamics under appropriate task constraints [2,54]. We therefore focus on the structure of the learned recurrent connectivity that gives rise to these dynamics.

Despite starting from unstructured random weights, learning consistently reshaped the recurrent matrix into a highly organized form, indicating that the task selects this structure rather than imposing it architecturally. When neurons were sorted by their peak activation time during the training sequence, the final recurrent connectivity exhibited a clear banded organization (Fig. 1D). Averaging weights along the diagonals revealed a Mexican-hat-like interaction profile characterized by strong local excitation and longer-range inhibition. Negative couplings reflect effective inhibition in the reduced model. Such profiles are canonical in attractor models and support low-dimensional ring manifolds capable of stable pattern completion [21,32–34,36,37,39,40,79].

Crucially, the learned connectivity was not purely symmetric. The eigenspectrum of the recurrent matrix, *W*_*hh*_, contained complex-valued modes (Fig. 1D, right), indicating the presence of antisymmetric structure. While most eigenvalues clustered near the origin, a small number of dominant modes emerged. In particular, the leading eigenvalue had a real component of 2.16 and a nonzero imaginary component of ± 0.15, indicating a dominant recurrent mode with weak rotational structure. Because these spectra are computed from the recurrent weight matrix *W*_*hh*_, this does not imply instability of the nonlinear dynamics. In a tanh RNN, the effective Jacobian is 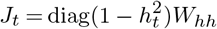, so the nonlinear gain contracts the spectrum during both replay and prediction. Further analysis of the effective gain can be found in Supplemental Material 6(d).

To isolate the contributions of these components, we decomposed the learned recurrent matrix into symmetric and antisymmetric parts (Fig. 1E-F). The symmetric component dominated the overall structure, as measured by its Frobenius norm (||*W*_*sym*_||_*F*_ = 3.65), which quantifies the total magnitude of the recurrent weights. It reproduced both the fully symmetric Mexican-hat trace profile and a purely real eigenspectrum (Fig. 1E), consistent with stable fixed-point and continuous attractor dynamics [21,36,37,79]. Although these dynamics resemble those of continuous attractor models, we do not claim that the trained network implements a mathematically exact continuous attractor. In contrast, the antisymmetric component, though smaller in weight magnitude (||*W*_*anti*_||_*F*_ = 1.94), accounted for approximately 35% of the total recurrent norm and exhibited imaginary eigenmodes within the unit circle (Fig. 1F), consistent with rotational or flow-like dynamics that bias activity propagation along the ring [53].

The spatial structure of the antisymmetric component closely resembles a noisy first spatial derivative of the symmetric Mexican-hat profile, with odd symmetry around zero offset and localization near the attractor peak (Fig. 1F, middle). Such derivative-like antisymmetric perturbations have been shown to convert static continuous attractors into dynamic solutions by introducing a controlled direction bias [53].

This decomposition shows how replay and prediction emerge from the learned connectivity. The symmetric component stabilizes representations on a low-dimensional attractor-like manifold, while the antisymmetric component breaks time-reversal symmetry and selects flow direction along that manifold. The resulting dynamics correspond to activity moving along a ring manifold, producing smooth phase advance in polar coordinates and coherent sequential structure during both replay and prediction (Fig. 1B). These trajectories do not perfectly overlap, indicating that the learned flow is flexible and not exactly aligned with the task manifold. The dominant outlier eigenmode combines strong symmetric gain with a weak antisymmetric contribution, suggesting that sequence generation arises from a mode that is gently rotated in state space rather than purely oscillatory dynamics [53,54,58,59].

These results demonstrate that optimization for one-step-ahead prediction from random connectivity reliably induces a mixed-symmetry recurrent organization, rather than converging to a purely symmetric attractor network. This provides evidence that symmetry breaking can be learned from task demands, while the residual mismatch between replay and prediction dynamics suggests that learning alone does not fully constrain the recurrent structure. These findings show that mixed-symmetry can emerge from task optimization alone, motivating a systematic examination of how structured recurrent inductive biases shape learning and performance.

### (b) CA3-like recurrent inductive bias improves replay, prediction, and learning dynamics

To assess how recurrent inductive bias shapes hippocampal-like computation, we compared networks initialized with different CA3-inspired recurrent motifs (identity, cyclic shift, Mexican-hat, see Methods section 2(b)) and evaluated autonomous replay and one-step-ahead prediction. Replay probes internally generated dynamics without informative input, whereas prediction tests whether learned temporal structure is maintained after withdrawal of external drive. Performance across both replay and prediction depended strongly on the initial recurrent structure. Networks initialized with a constructed Difference-of-Gaussians Mexican-hat (center-surround) connectivity (see Methods section 2(e)) consistently achieved the strongest performance across all metrics (Tables 1-2; Fig. 2).

#### (i) Replay

During replay, Mexican-hat initialization with an initial symmetry value of *α*_0_ = 0.70 yielded the highest ring-decode *R*^2^ and near-maximal residual lag-1 autocorrelation (Table 1), indicating that internally generated activity remains confined to the learned ring manifold while evolving smoothly over time. More information on metrics used can be found in the Supplemental Material section 6(e).

Random initialization also supported replay, showing that task-driven learning alone can induce continuous-attractor-like dynamics. However, replay trajectories were less consistent across runs and showed weaker alignment between replay and prediction than in Mexican-hat networks (Fig. 2D). In contrast, identity initialization, which is maximally symmetric and lacking lateral interactions, failed to support replay as shown by exhibiting low ring-decode *R*^2^ and poor temporal coherence. Thus, temporal persistence alone is insufficient for autonomous sequence generation. Further results from the identity and cyclic shift matrices can be found in the Supplemental Material section 6(c).

#### (ii) Prediction

Prediction performance further separated the connectivity families (Table 2). Mexican-hat networks simultaneously maximized angular accuracy (ring-decode *R*^2^ and mean resultant length *R*) while minimizing mean squared error. Cyclic shift connectivity, which hardwires directionality, improved prediction and replay relative to identity initialization but underperformed Mexican-hat connectivity and exhibited replay breakdown at higher symmetry levels (Supplementary Figs. S4-S5). Thus, directionality alone is also insufficient without a stabilizing recurrent scaffold.

Structured inductive bias also altered optimization dynamics. The constructed Mexican-hat initialization often began at lower loss and exhibited a pronounced early reduction in error compared to random connectivity (Fig. 2A,B). For specific initial symmetry values, training showed a two-phase learning profile that was absent under random initialization, with an early rapid loss decrease followed by slower refinement [80,81]. The two-phase learning dynamics suggest that networks first learn the input-output mapping and subsequently refine the recurrent structure that stabilizes and directs activity along the task manifold. This indicates that structured recurrence not only improves final performance but also places the network in a more favorable regime for learning. These advantages raise the question of what recurrent structure is ultimately selected by optimization under different initial conditions. We therefore next examine how learning reshapes recurrent connectivity and whether it converges to a common dynamical regime across structured initializations.

### (c) Learning converges to a mixed-symmetry regime supporting sequential dynamics

Across structured initializations, learning consistently drove recurrent connectivity toward an intermediate balance between symmetric and antisymmetric components, quantified by the symmetry index *α*(*t*) (Fig. 2C). Despite wide variation in initial symmetry, *α*(*t*) converged to a predominantly symmetric regime with a persistent antisymmetric component, indicating that intermediate symmetry is task-selected rather than incidental. Across all conditions, the best performance was obtained for off-centered Mexican-hat connectivity initialized with an intermediate symmetry level (*α*_0_ = 0.70), which places the network near the task-selected symmetry regime from the outset (Fig. 2A-C).

The learned recurrent weights preserve a Mexican-hat-like center-surround profile characterized by strong local excitation and broader inhibition (Fig. 2E-F). Decomposition of the final connectivity revealed that the symmetric component dominated the overall structure and eigenspectrum (Fig. 2G, ||*W*_*sym*_||_*F*_ = 3.7), consistent with stabilization of an attractor manifold. In contrast, the antisymmetric component was smaller in magnitude (Fig. 2H; ||*W*_*anti*_||_*F*_ = 0.42, ||*A*||*/*(||*S*||+||*A*||) = 0.10) but introduced imaginary eigenmodes that bias activity propagation. As in the randomly initialized networks, the learned antisymmetric component closely resembles a noisy spatial derivative of the symmetric Mexican-hat profile, with odd symmetry around zero and localization near the attractor peak (Fig. 2H, middle). This derivative-like structure is consistent with theoretical predictions that weak antisymmetric perturbations of attractor-like manifolds induce directional drift without destabilizing the underlying manifold [53].

Functionally, this mixed-symmetry regime supports dynamics that are neither purely static nor purely feedforward. Replay and prediction trajectories were more closely aligned for Mexican-hat initializations than for random connectivity (Fig. 2D), indicating that internally generated dynamics reflect the learned predictive structure. Spectral analyses further showed that the best-performing initializations already contained weak imaginary components (Fig. 2E), which were preserved and refined during training (Fig. 2F-H).

These results show that task optimization reliably selects an intermediate symmetry regime in which symmetric recurrence stabilizes representations while structured antisymmetry enables controlled temporal progression. Task learning can recover mixed-symmetry recurrent structure from random connectivity, but structured inductive bias accelerates learning, stabilizes dynamics, and yields the most robust replay and prediction. In the next section, we examine how a biologically plausible plasticity mechanism, specifically a modified STDP rule, can give rise to this structured symmetry breaking.

### (d) Heterogeneous STDP induces symmetry breaking in Mexican-hat connectivity

To connect the mixed-symmetry structure observed in the trained RNNs to biologically plausible learning mechanisms, we analyzed the RNN structure emergent from a modified STDP rule that interpolates between antisymmetric and symmetric plasticity. This modified STDP rule replaces backpropagation used in the previous experimental setups. Classical STDP is temporally asymmetric, strengthening synapses when presynaptic activity precedes postsynaptic firing and weakening the reverse. While such causal learning supports sequence formation, it does not by itself produce the stabilized center-surround structure observed in the learned recurrent connectivity (Figure 2). Conversely, purely symmetric Hebbian plasticity produces stable attractors but lacks directional bias [36].

Under a traveling Gaussian input, the modified STDP learning rule produces a family of synaptic weights whose spatial interaction pattern is determined by the symmetry parameter *ϵ* (see Methods section 2(f) and analytical derivation in Supplementary section 6(f); Fig. 3). For *ϵ* = 0, the rule reduces to standard STDP and yields a strongly antisymmetric interaction profile, dominated by imaginary eigenvalues and fast drift of the activity bump (Fig. 3A). For *ϵ* = 1, a symmetric Mexican-hat profile is produced with a purely real eigenspectrum (Fig. 3B). For intermediate values (*ϵ* = 0.85), the learned synaptic weight profile becomes a weakly tilted Mexican hat, preserving the stabilizing center-surround structure while introducing a small antisymmetric component (Fig. 3C). The corresponding eigenspectrum contains dominant real modes with only small imaginary components, yielding a stable activity bump that propagates at a constant velocity.

Although *ϵ* plays a role analogous to the symmetry index *α*, the two are not identical. *ϵ* parametrizes symmetry at the level of synaptic plasticity, whereas *α* quantifies the symmetry of the overall connectivity. Nevertheless, intermediate values of *ϵ* naturally give rise to recurrent connectivity in the same mixed-symmetry regime favored by task optimization in the RNNs.

The analysis of the modified STDP learning rule shows that the flow-stabilized attractor regime is not merely an outcome of backpropagation but is also accessible to biological circuits via simple tuning of the retrograde plasticity signal (*ϵ*). In biological terms, *ϵ* may correspond to the strength of retroactive potentiation observed in recent STDP experiments in CA3 [69] allowing the hippocampus to interpolate between storage (attractor) and replay (flow) modes through purely local changes.

## 4. Discussion

### (a) Symmetry breaking as a mechanism for hippocampal sequence generation

We have built on previous work to provide a quantitative analysis of partial symmetry breaking in recurrent connectivity as a circuit-level principle that enables hippocampal networks to jointly support stable memory representations, internally generated replay, and predictive sequence dynamics. Using recurrent neural networks trained on a prediction task, we show that these functions emerge most robustly in an intermediate regime dominated by symmetric structure but augmented by a weaker antisymmetric component. This mixed-symmetry organization stabilizes internal representations while enabling directed flow along them, yielding dynamics that unify replay and prediction within a single recurrent circuit.

This regime is not imposed but selected by learning. Across random and structured initializations, networks reliably converge toward a tilted Mexican-hat-like connectivity, suggesting that symmetry breaking acts as a low-dimensional control parameter governing the balance between stability and motion. These findings support a view in which hippocampal circuitry is shaped by biological inductive biases that constrain learning toward a family of dynamical regimes well suited for sequential memory, with experience refining task-specific operating points within that space rather than discovering structure *de novo*.

Symmetric recurrent connectivity stabilizes attractor states but suppresses temporal structure, whereas strong asymmetry introduces directionality at the expense of stability. A long line of theoretical work has shown that symmetry breaking in recurrent networks is essential for generating ordered temporal dynamics [13,20,41,53,61]. In hippocampal models, asymmetric interactions have been invoked to explain path integration, sequential recall, and internally generated trajectories [14,21,54,58,60].

Our results extend this framework by showing that replay and prediction emerge most robustly when symmetric and antisymmetric components coexist in a controlled ratio. Networks trained from various initial conditions reliably converge toward this mixed-symmetry regime, suggesting that partial symmetry breaking is a task-selected dynamical solution.

### (b) Circulant connectivity and flow-equivariant dynamics in hippocampal networks

An important theoretical insight emerging from this work is the role of translation-invariant recurrent structure, formalized as Toeplitz or, more specifically, circulant connectivity [55,82]. In such networks, synaptic weights depend primarily on relative position, yielding spatial Fourier eigenmodes. Symmetric Toeplitz structure selectively stabilizes low-frequency modes, producing smooth continuous attractors, whereas antisymmetric components introduce imaginary eigenvalues that generate coherent drift along the attractor manifold.

From this perspective, the balance between symmetric and antisymmetric structure directly determines whether dominant modes are stationary or propagating. Our findings show that learning consistently drives recurrent connectivity toward this structured regime, even when starting from random matrices. This suggests that circulant-like structure is not merely analytically convenient but an efficient solution for predictive sequence learning.

Approximate translation invariance is biologically plausible in hippocampal CA3, where recurrent connectivity exists and lacks obvious spatial anchoring. Experience-dependent plasticity would naturally reinforce relative rather than absolute coding, while deviations from perfect symmetry provide a principled mechanism for generating directed flow without disrupting representational stability. In this way, partial symmetry breaking transforms a static attractor into a flow-stabilized representation.

Our model provides a concrete instantiation of this principle: partial symmetry breaking within a circulant recurrent scaffold yields dynamics that unify replay and prediction through flow along an internal representational manifold. In this sense, hippocampal recurrent dynamics implement a form of flow-equivariant computation [82], in which task-relevant structure is preserved under continuous transformations while temporal progression is encoded through controlled motion along that structure.

### (c) Relationship to prior models

Zhang’s seminal work explicitly decomposed recurrent connectivity into even (symmetric) and odd (antisymmetric) components, and demonstrated that the odd component generates rotational flow in head-direction networks [53]. He further showed that optimal dynamics arise in an intermediate regime dominated by symmetry but containing a weaker antisymmetric component, and that the antisymmetric structure approximates the spatial derivative of the symmetric component.

Our results closely parallel this model but extend it in several key ways. First, whereas Zhang prescribed the relative strength of even and odd components to implement velocity integration, we show that this balance can emerge through learning under a predictive objective. Second, Zhang’s model relies on externally driven velocity inputs, whereas our networks generate replay and prediction internally, in the absence of explicit motion signals. Finally, although the learned antisymmetric component closely resembles the spatial derivative of the symmetric Mexican-hat profile, this relationship arises naturally from optimization rather than being imposed. Together, these findings extend symmetry-based theories from hand-designed integrators to learned hippocampal dynamics.

These results can be viewed as a continuous generalization of earlier fast-slow attractor models [13,14,43–45], in which stabilization and transition dynamics were implemented as separate processes, showing instead that both can emerge within a single recurrent circuit through learned partial symmetry breaking.

Recent theoretical syntheses emphasize that neural circuits often operate between pure attractor and pure integrator regimes [83]. Our results identify partial symmetry breaking as a concrete mechanism for realizing such intermediate dynamics in hippocampal networks.

### (d) Structured versus random recurrent connectivity

Random recurrent networks can exhibit rich dynamics, particularly near the edge of chaos [57], and training can harness this richness for computation [75,84]. However, such networks often require fine-tuning to suppress instability and may lack robustness across tasks.

In contrast, we find that structured inductive biases, specifically Mexican-hat-like connectivity with partial symmetry breaking, provide a stable substrate for learning. Networks initialized near this regime learn faster, converge more reliably, and exhibit more interpretable dynamics than those initialized randomly. This aligns with work showing that population structure constrains and stabilizes computation in recurrent circuits [74,85]. From a biological standpoint, these results support the idea that hippocampal circuitry may be developmentally biased toward certain connectivity motifs, with learning refining rather than discovering them *de novo*.

### (e) Biological plausibility

The coexistence of symmetric and asymmetric connectivity need not imply multiple specialized learning rules. Synapses exhibit substantial heterogeneity in timing, plasticity, and short-term dynamics [58,69,71,86]. Our heterogeneous STDP results demonstrate that realistic synaptic diversity can naturally give rise to the structured symmetry breaking required for flow, providing a plausible biological substrate for the dynamics observed here.

The relationship between firing-rate attractor models and biologically realistic spiking networks is an important consideration. CA3 pyramidal neurons fire sparsely, raising the question of how smooth attractor dynamics emerge from low-rate, temporally discrete spike trains. The present model does not attempt to directly resolve this gap, instead adopting a mesoscopic, population-rate description in which recurrent symmetry breaking shapes effective dynamical flow. However, prior work has shown that trained rate networks can be mapped to spiking implementations that preserve key dynamical features [35]. Consistent with this, applying this mapping to the present model yields replay and theta phase precession in a network of leaky integrate-and-fire neurons [2], suggesting that the learned dynamical structure can, in some cases, be expressed in a spiking regime. That said, whether this specific flow-stabilized attractor can be robustly implemented in sparse spiking networks remains an important direction for future work.

The Mexican-hat profiles reported here represent effective interaction at the level of population dynamics. In biological CA3 circuits, a comparable center-surround structure would arise through excitatory-inhibitory microcircuit organization. Several studies have demonstrated that recurrent networks respecting Dale’s principle can reproduce similar dynamical regimes by explicitly separating excitatory and inhibitory populations [87–91]. The present study focuses on the dynamical consequences of symmetry structure in the effective recurrent operator, leaving the detailed microcircuit implementation to future work.

Experimental measurements suggest that CA3 recurrent excitatory connectivity is sparse and only weakly distance-dependence across several species [72,77,92–94]. The Mexican-hat profile used here should therefore be interpreted as an effective population-level connectivity motif rather than a direct anatomical description. In hippocampal circuits, structured inhibition, heterogeneous plasticity rules, and network dynamics can shape the interactions experienced by pyramidal populations, potentially giving rise to center-surround structure at the population level despite sparse underlying connectivity. An important direction for future work is to determine whether the mixed-symmetry regime identified here persists in biologically constrained architectures incorporating sparse connectivity and Dale’s principle.

An important distinction is that the traveling waves observed in our model are representational rather than explicitly anatomical. However, these two forms of dynamics may be related. Anatomical traveling waves have been observed along the septotemporal (dorsal-ventral) axis of the hippocampus, demonstrating that activity can propagate across this axis in a coordinated manner [19,66]. At the same time, place fields are spatially distributed and partially redundant, such that neurons representing nearby or even similar locations are not arranged topographically but instead interspersed across the network [8,95–98]. In such a distributed code, a propagating anatomical wave need not correspond to a contiguous trajectory in physical space. Instead, as activity moves along the dorsal-ventral axis, it could sequentially recruit neurons whose place fields collectively form a trajectory through the environment, even though those neurons are spatially interspersed within the hippocampus. Thus, the structured propagation required for anatomical traveling waves could arise from an underlying recurrent connectivity motif with local excitation and longer-range inhibition, qualitatively consistent with a tilted Mexican-hat profile. We therefore speculate that the same class of connectivity that supports anatomical wave propagation may also give rise to the internally generated representational sequences observed during replay and prediction. Importantly, this interpretation does not imply that anatomical waves directly encode spatial trajectories, but rather that they reflect a more general dynamical principle that structured recurrent interactions can generate propagating activity patterns which, when combined with a distributed spatial code, appear as flexible sequences in representational space. This hypothesis makes a testable prediction that, if correct, connectomic analyses of CA3 should reveal effective interactions consistent with a weakly asymmetric (tilted) center-surround structure along the septotemporal axis.

Our findings support a view of hippocampal CA3 as a flow-stabilized recurrent circuit in which memory, replay, and prediction emerge from a balance between symmetric stabilization and directed flow. Conceptual frameworks emphasize that the hippocampus is fundamentally a sequence generator, with internally generated, ordered dynamics forming the basis of memory and prediction rather than static representations [3,12,13,19,22,31,99]. We find that recurrent networks trained on predictive sequence tasks converge to a regime dominated by Mexican-hat-like symmetric connectivity with a smaller but essential antisymmetric component. This mixed-symmetry organization is consistent with experimental observations in CA3, where recurrent synapses exhibit largely symmetric plasticity optimized for autoassociative storage [69], alongside structural and functional asymmetries between pyramidal subtypes that support sequential activation [71–73,77]. The CA3 is difficult to directly measure connectivity, but more generally, connectomic studies show strongly directed and largely acyclic connectivity in cortical microcircuits, indicating biological networks can display structural asymmetry [100]. Although learning from random connectivity can recover this regime, structured CA3-like inductive biases accelerate convergence and stabilize dynamics. This suggests that hippocampal circuitry constrains learning toward dynamical regimes well suited for sequential memory tasks.

This framework yields clear experimental predictions. Selectively perturbing directional coupling should impair replay speed or directionality without abolishing completion (e.g. targeted optogenetic manipulation of CA3 ensembles [101], while state-dependent modulation of CA3 input pathways should bias internal flow without disrupting the underlying attractor geometry [86]. While the exact symmetry level, propagation speed, and gain depend on task demands, the principle of dominant symmetry augmented by weak derivative-like antisymmetry appears to be task-general. These findings argue that hippocampal sequence generation reflects an interaction between biological inductive biases and task-driven learning, rather than learning alone.

These results suggest that hippocampal replay and prediction emerge from a delicate balance between stability and flow, implemented through partial symmetry breaking in recurrent connectivity. Circulant structure defines the internal representational manifold and antisymmetry determines how that manifold is traversed. Flow is therefore a functional requirement for hippocampal computation, enabling internal models to be dynamically explore rather than statically recalled. More broadly, this work suggest that CA3-like inductive bias does not prescribe a specific computation, but instead constrains learning toward a family of dynamical regimes well suited for sequential memory tasks—a tilted Mexican hat. Structured connectivity restricts the search space of learning, while task demands select a particular operating point within that space. The convergence of mixed-symmetry structure across random initialization, structured bias, and biologically plausible plasticity argues that this organization is not a task artifact. Although the precise symmetry level, propagation timescale, and antisymmetric gain depend on task demands, the organizing principle of dominant symmetry combined with weak, derivative-like antisymmetry appears to be general.

## 5. Conclusion

Several limitations point to important directions for future work. We focused on a single prediction task and abstracted away cell-type diversity, neuromodulation, and short-term synaptic dynamics, all of which are known to influence hippocampal computation. Whether the same mixed-symmetry regime generalizes to non-spatial memories, branching sequences, or goal-dependent dynamics remains an open question. Extending this idea to multi-area models incorporating CA1, entorhinal cortex, and state-dependent modulation will be essential for linking circuit dynamics more directly to behavior.

The framework developed here yields clear experimental predictions. Selective perturbations of directional coupling within CA3 should impair replay speed or directionality without abolishing pattern completion, while state-dependent modulation of CA3 inputs should bias internal flow while preserving the underlying attractor geometry. Advances in large-scale electrophysiology and imaging now make it possible to directly test whether hippocampal population dynamics exhibit the predicted weak rotational components associated with partial symmetry breaking.

In conclusion, these results support a view of hippocampal CA3 as a flow-stabilized recurrent circuit in which memory, replay, and prediction arise from the interaction between inductive biases and experience-dependent learning. Partial symmetry breaking provides a principled mechanism for balancing stability and motion, offering a unifying circuit-level account of hippocampal sequence generation across behavioral states.

## Acknowledgements

This research was supported by an NIH Director’s Pioneer Award (DP1NS149613) and by a grant from the Office of Naval Research (N00014-23-1-2069).

## 6. Supplemental Material

### (a) Pretraining as evidence for structural inductive bias

**Figure S1:**
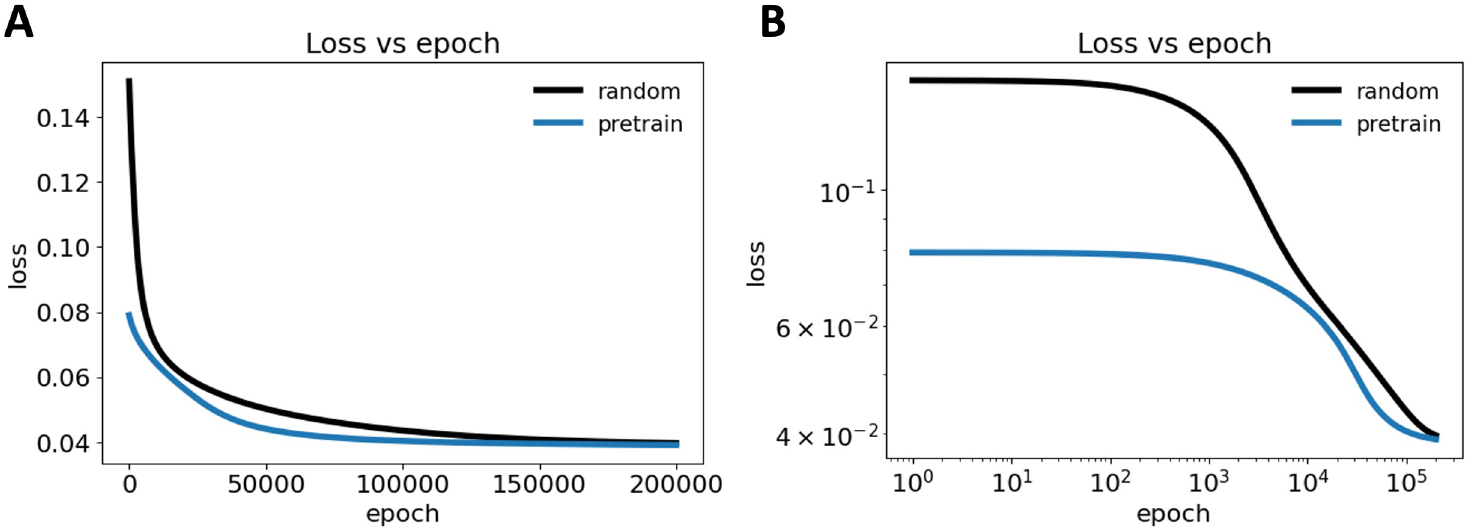
Effect of pretraining on learning dynamics. **(A)** Training loss as a function of epoch shown on linear axes. Networks initialized with pretrained recurrent weights (blue), obtained from the final weights of the randomly initialized network, begin at a lower loss and exhibit faster early optimization compared to networks trained from random initialization (black). **(B)** Same data plotted on log-log axes, highlighting differences in early and intermediate training regimes. Pretrained networks avoid the prolonged high-loss plateau observed under random initialization and transition earlier into the rapid loss-decrease regime. In both cases, training converges to a similar final loss.

Pretraining the recurrent connectivity using the final weights obtained from a randomly initialized network significantly alters the early learning dynamics without changing the eventual performance ceiling (Fig. S1). As shown in both linear (Fig. S1A) and log-log (Fig. S1B) axes, pretrained networks start at a markedly lower loss and bypass the extended high-loss plateau characteristic of random initialization. This leads to faster progress through early and intermediate training epochs. Despite these differences in learning trajectory, both conditions converge to nearly identical final loss values, indicating that pretraining primarily improves optimization efficiency rather than the final representational capacity of the model. These results suggest that structure acquired during prior training places the network in a favorable regime of parameter space, facilitating faster learning while reserving the same asymptotic solution.

### (b) Model hyperparameters

**Table S1:** Model hyperparameters.

| hyperparameter | value |
| --- | --- |
| number of units | 100 |
| number of epochs | 100,000 |
| sequence length | 100 |
| learning rate | 0.01 |

### (c) Identity and cyclic shift initializations: isolating persistence, directionality, and their limits in CA3-like circuits

To further dissect which structural features of CA3 recurrent connectivity are necessary for hippocampal-like replay and prediction, we examined two extreme and deliberately simplified structures: identity and cyclic shift connectivity. These architectures isolate the complementary computational primitives of temporal persistence and directional propagation (symmetric and antisymmetric aspects) that are each present in CA3, although not in isolation.

The identity matrix represents a maximally symmetric, self-coupled network in which each unit predominantly feeds back onto itself. We include this case as a conceptual control that isolates temporal persistence without lateral interactions. Such a network can maintain activity locally and support short-horizon, input-driven prediction. However, it fundamentally fails to capture CA3’s role as an associative recurrent circuit. Identity-initialized networks reliably learned one-step prediction when driven by external input but failed at autonomous replay. The resulting weight profiles remained identity-dominated with only weak lateral structure, which was insufficient to sustain a self-organized sequence when driven by noise. Thus, persistence alone is not sufficient for sequential dynamics or replay. This reinforces the idea that CA3 replay requires not just stability, but structured interactions that organize population activity along a low-dimensional manifold.

In contrast, cyclic shift connectivity (Fig. **??**) imposes a purely directional recurrent structure, in which each neuron excites its success along a ring. This architecture hardwires a translation operator into the recurrent matrix and naturally produces a sequential propagation of activity. As such, it captures the CA3 function of directional progression through a learned sequence and is consistent with experimental and theoretical work implicating antisymmetric or non-reciprocal connectivity arising from temporally antisymmetric plasticity such as STDP. Indeed, antisymmetric components in CA3 may reflect experience-dependent strengthening of forward transitions during repeated trajectories, theta sequences, or sharp-wave ripple replay.

**Figure S2:**
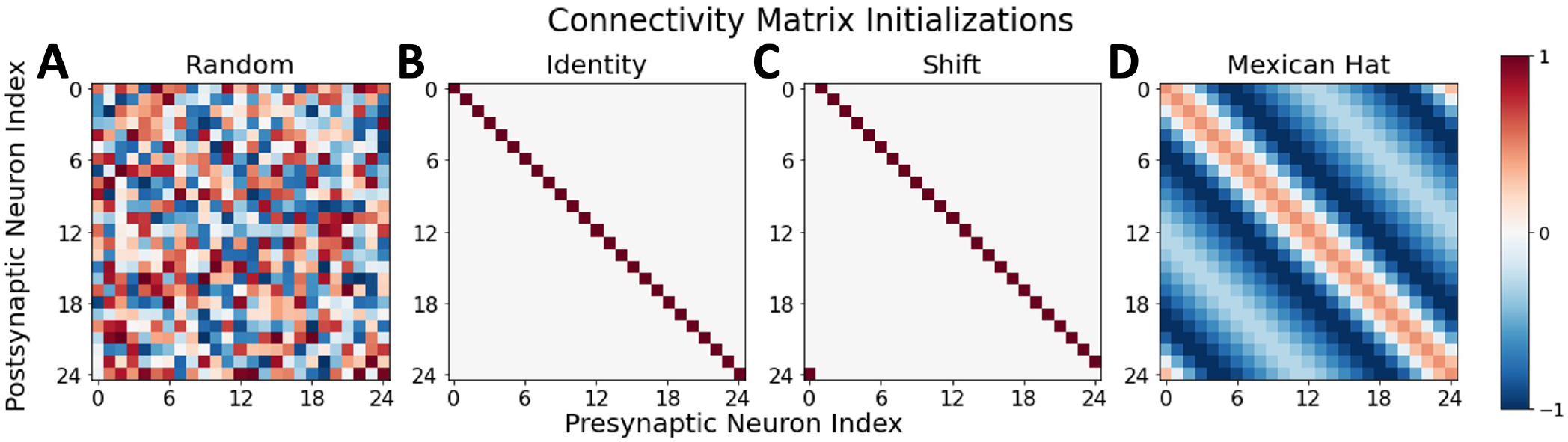
Connectivity matrix initialization families. Visualization of the four recurrent families used in this study: **(A)** random connectivity, with weights sampled from a zero-mean distribution (PyTorch baseline). **(B)** identity connectivity, in which each neuron connects only to itself. **(C)** cyclic shift connectivity, where each neuron projects to its immediate successor on a ring (with wrap-around), implementing a discrete translation operator. **(D)** Mexican-hat connectivity, constructed as a difference-of-Gaussians kernel on a ring, producing local excitation and broader surround inhibition. Identity and cyclic shift connectivity represent limiting cases of purely symmetric (self-coupled) and purely directional (translation-only) structure, respectively, while Mexican-hat connectivity combines structured excitation and inhibition to support attractor dynamics.

**Figure S3:**
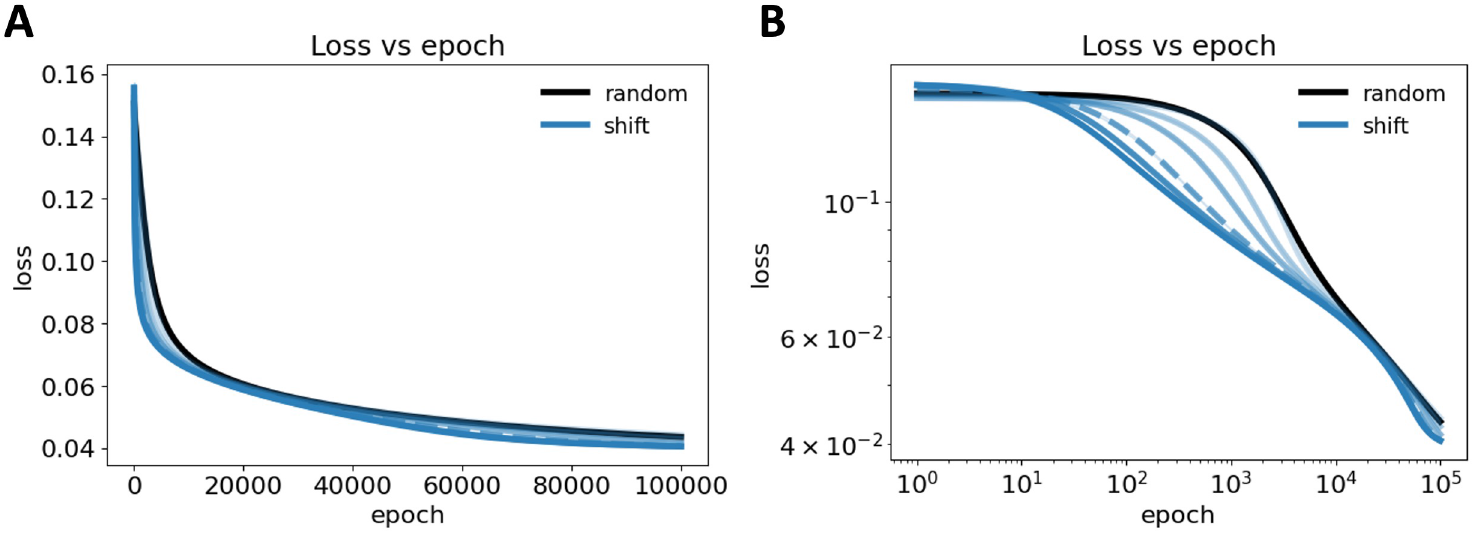
Training loss for cyclic shift-initialized recurrent networks across symmetry levels. Training loss as a function of epoch for cyclic shift initializations (blue) spanning initial symmetry ratios of *α*_0_ ∈ [0, 1] compared to random initialization (black). Light blue is the most antisymmetric and the symmetry ratio increases as the blue line darkens. The dashed blue curve is *α*_0_ = 0.60, the beginning of the regime where replay breaks down despite comparable training loss. **(A)** linear scale and **(B)** log-log scale. Overall, cyclic shift-initialized networks exhibit similar final and initial losses to random connectivity, while early training dynamics are altered. Stable replay here is not guaranteed.

However, pure cyclic shift connectivity lacks the bidirectional, stabilizing interactions required for attractor-like memory retrieval. Our results make this limitation explicit. While cyclic shift-initialized networks were able to outperform identity networks on both replay and prediction, their ability to support replay depended critically on the initial symmetry-antisymmetry balance. As the initial symmetry ratio *α*_0_ increased, the structure became more similar to the identity matrix, with a complete sharp breakdown in replay behavior at *α*_0_ = 0.60 indicated by ring-decode *R*^2^ collapse, residual temporal coherence deterioration, and polar trajectories which degenerated into fixed or noisy states (Fig. S4-S5) Indeed, at *α*_0_ = 0, the recurrent matrix is a pure cyclic shift matrix, but at *α*_0_ = 1, the matrix has the same limitations as the identity matrix. Notably, prediction performance remained largely intact across this transition, indicating that locally accurate state updates can still be learned even when the global autonomous dynamics fail.

This highlights a key computational distinction. Prediction can be supported by a locally correct transition operator under external drive, whereas replay requires a globally consistent dynamical system with both a stable manifold and a directed flow along it. Pure cyclic shift connectivity provides direction without sufficient stabilization; identity provides stabilization without direction. Random connectivity can, through learning, self-organize both to some degree but does so less intuitively than biologically inspired structures.

**Figure S4:**
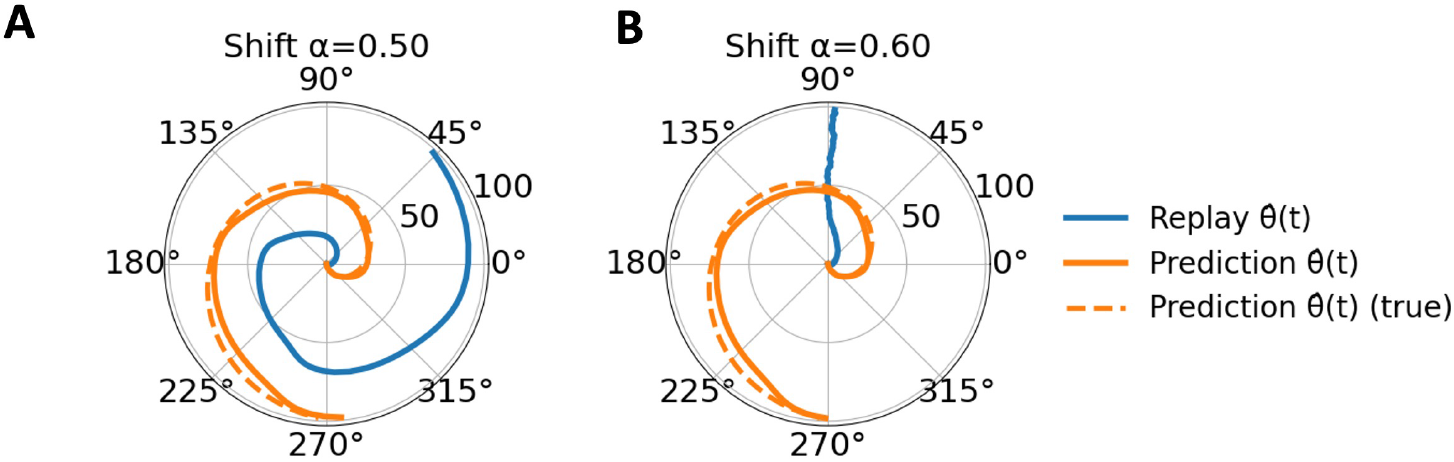
Breakdown of replay for cyclic shift-initialized networks at higher symmetry. Polar trajectories of learned replay (blue) and prediction (orange; dashed indicated ground truth) for cyclic shift-initialized recurrent connectivity with **(A)** *α*_0_ = 0.50 and **(B)** *α*_0_ = 0.60. At *α*_0_ = 0.50, replay follows a coherent progression around the ring, whereas increasing symmetry to *α*_0_ = 0.60 leads to a collapse of replay dynamics despite preserved prediction performance, indicating a sharp transition in the ability of cyclic shift connectivity to support stable autonomous sequence generation.

**Figure S5:**
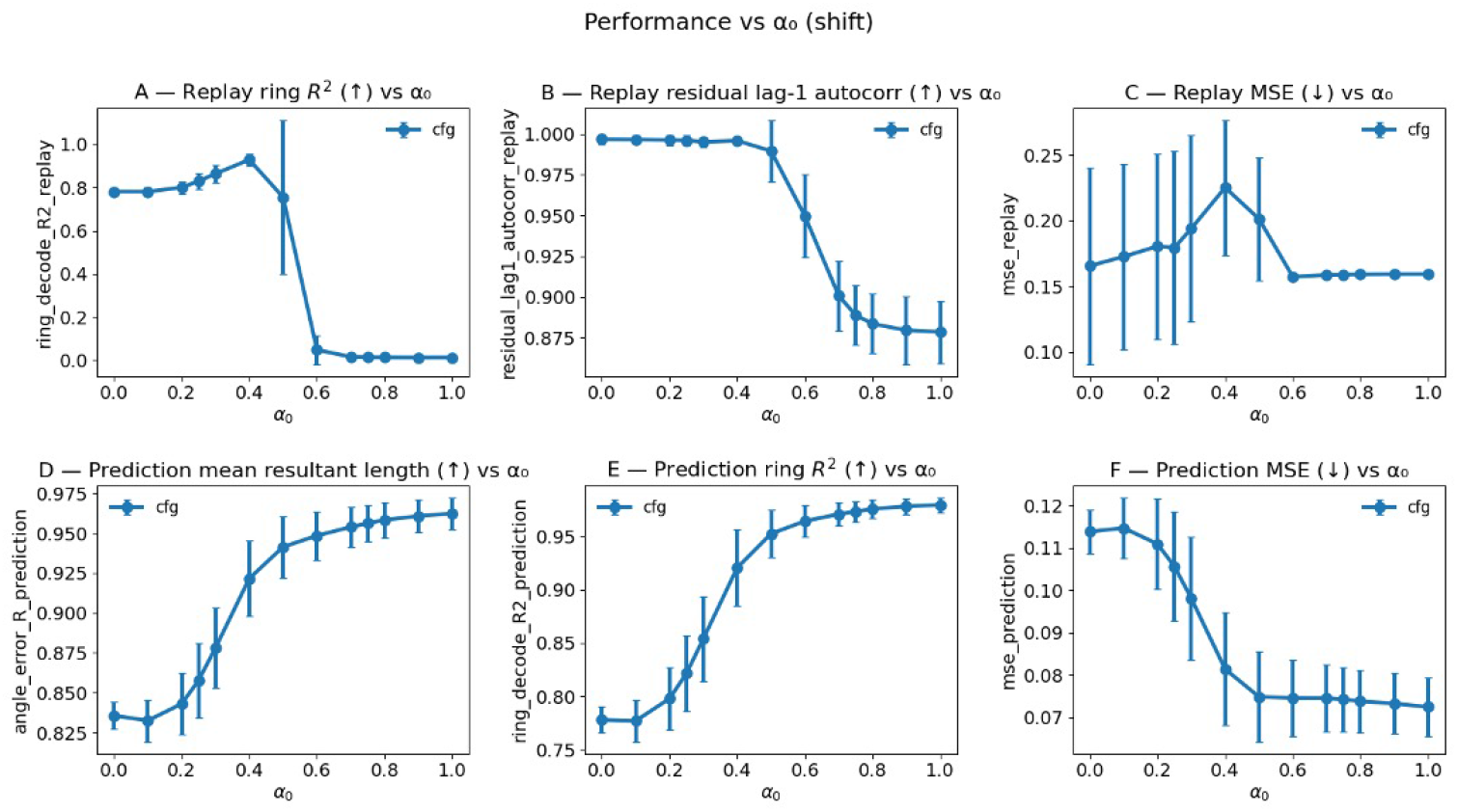
Performance as a function of initial symmetry for cyclic shift connectivity. Replay and prediction performance metrics plotted as a function of the initial symmetry ratio *α*_0_ for cyclic shift-initialized recurrent connectivity. Top row: replay metrics including ring-decode *R*^2^ **(A)**, residual lag-1 autocorrelation **(B)**, and replay MSE **(C)**. Bottom row: prediction metrics including mean resultant length of angular error **(D)**, ring-decode *R*^2^ **(E)**, and prediction MSE **(F)**. Replay performance exhibits a sharp breakdown at *α*_0_ = 0.60, marked by the dramatic drop in ring-decode *R*^2^ and reduced temporal coherence despite increased prediction performance. This difference indicates that increasing symmetry in cyclic shift connectivity impairs autonomous replay dynamics while leaving input-driven prediction intact.

Across structured initializations, training consistently drove the recurrent connectivity toward an intermediate symmetry regime, quantified by convergence of *α*(*t*) (Fig. S6). Fully symmetric networks (i.e. identity) rapidly gained antisymmetric structure. Cyclic shift networks, spanning fully antisymmetric to symmetric initial conditions, converged toward a mid-range *α* value. Of note, the intermediate range here is lower than that of Mexican-hat initialization. This is likely connected to the breakdown in replay behavior observed at *α*_0_ = 0.60. This convergence is again not an artifact of initialization, but instead an emergent, task-selected property, reinforcing the conclusion that hippocampal-like computation requires a balance between symmetric stabilization and antisymmetric flow.

A tilted Mexican-hat connectivity obtains this balance naturally. By providing a center-surround attractor scaffold from the outset, Mexican-hat initialization supports both stable memory representations and controlled propagation, yielding superior replay fidelity, tighter alignment between replay and prediction than either cyclic shift or identity connectivity. In this context, the cyclic shift and identity initializations serve as informative boundary cases that reveal that neither directionality nor persistence alone is sufficient. Instead, CA3-like replay emerges most robustly when directional biases are embedded with a stabilizing recurrent architecture. This is a regime that learning reliably discovers, but that structured inductive bias makes easier to reach and clearer to interpret.

**Figure S6:**
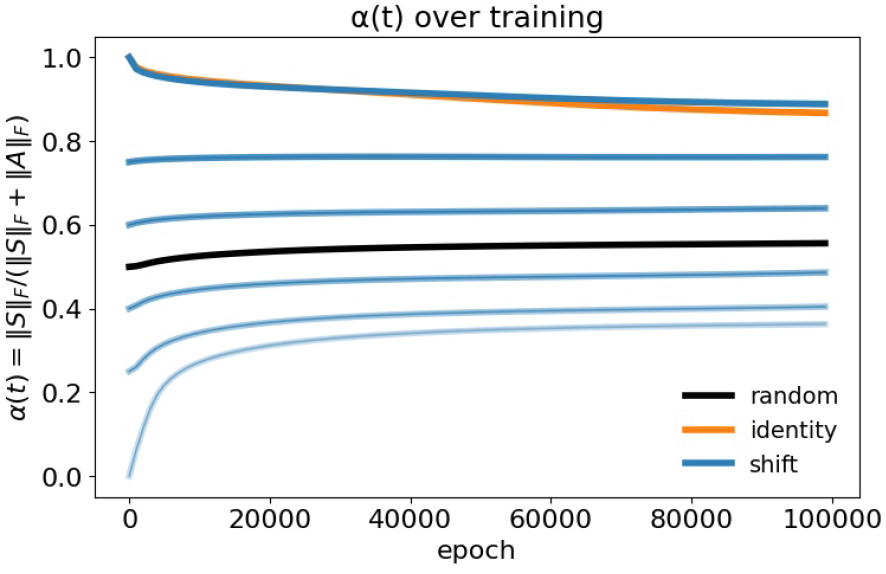
Training drives recurrent connectivity toward an intermediate symmetry regime. Evolution of the symmetry index *α*(*t*) = ||*S*||_*F*_ */*(||*S*||_*F*_ +||*A*||_*F*_) during training for networks initialized with random (black), identity (orange), and cyclic shift (blue; multiple initial *α*_0_) recurrent connectivity. The initial *α* value, *α*_0_, is given by the value at epoch 0. Across initial conditions, learning consistently drives *α*(*t*) toward an intermediate range, indicating convergence to a mixed symmetric-antisymmetric recurrent regime.

Together, these results argue that CA3 is best understood not as a pure attractor (identity) network or a feedforward sequence generator (pure cyclic shift), but as a flow-stabilized recurrent system in which symmetric and antisymmetric interactions are jointly essential. Identity and cyclic shift initializations clarify the computational roles of these components, while Mexican-hat connectivity demonstrates how their integration supports replay and prediction in a single recurrent circuit.

### (d) Effective Jacobian and gain statistics

In an RNN with tanh nonlinearity, stability is determined by the instantaneous Jacobian

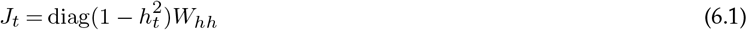

where *h*_*t*_ denotes the hidden state.

Because the network operates in a driven regime without a single fixed point, the relevant operating regime is defined by trajectories during replay and prediction. We therefore quantified the effective gain of the tanh nonlinearity along these trajectories. The tanh derivative acts as a state-dependent gain that rescales the recurrent matrix, determining the effective dynamics.

Across runs with random initialization, the mean derivative during replay was 0.31 ± 0.005 (median 0.14), with approximately 43% of activations having derivative less than 0.1, indicating substantial saturation (see Fig. S7 for full distribution). Similar values were observed during prediction (mean ≈ 0.27). Networks initialized with Mexican-hat connectivity exhibited slightly lower gains (mean 0.23-0.27), consistent with a more saturated operating regime.

These results imply that the effective Jacobian is a state-dependent, diagonally scaled version of *W*_*hh*_, which contracts the spectrum relative to the raw recurrent matrix. Consequently, eigenvalues of *W*_*hh*_ exceeding unity do not imply instability of the nonlinear system. Instead, the eigenspectrum of *W*_*hh*_ should be interpreted as reflecting the dominant recurrent interaction structure, with the nonlinear gain determining the effective dynamical regime.

**Table S2:** Gain analysis.

| Condition | Mean gain | Median gain | Percent < 0.1 |
| --- | --- | --- | --- |
| Random replay | $0.305 \pm 0.005$ | .0142 | 43% |
| Random prediction | $0.272 \pm 0.002$ | 0.116 | 47% |
| Mexican hat replay | $0.268 \pm 0.005$ | 0.083 | 53% |
| Mexican hat prediction | $0.228 \pm 0.003$ | 0.062 | 58% |

The broad distribution of tanh gain indicates that the network operates in a heterogeneous dynamical regime rather than a globally linear one. Many units are partially saturated, which is consistent with recurrent stabilization, while a smaller subset remains in a higher-gain regime capable of supporting directional state transitions. This is qualitatively consistent with the connectivity decomposition into a dominant symmetric scaffold plus a weaker antisymmetric component.

### (e) Metrics computed

#### (i) Ring decoding via cosine-sine regression

We evaluate decoding of a circular variable using linear (ridge) regression on cosine and sine targets, reporting performance via *R*^2^ [102]. Decoding cos *θ* and sin *θ* avoids angular wrap-around discontinuities.

**Figure S7:**
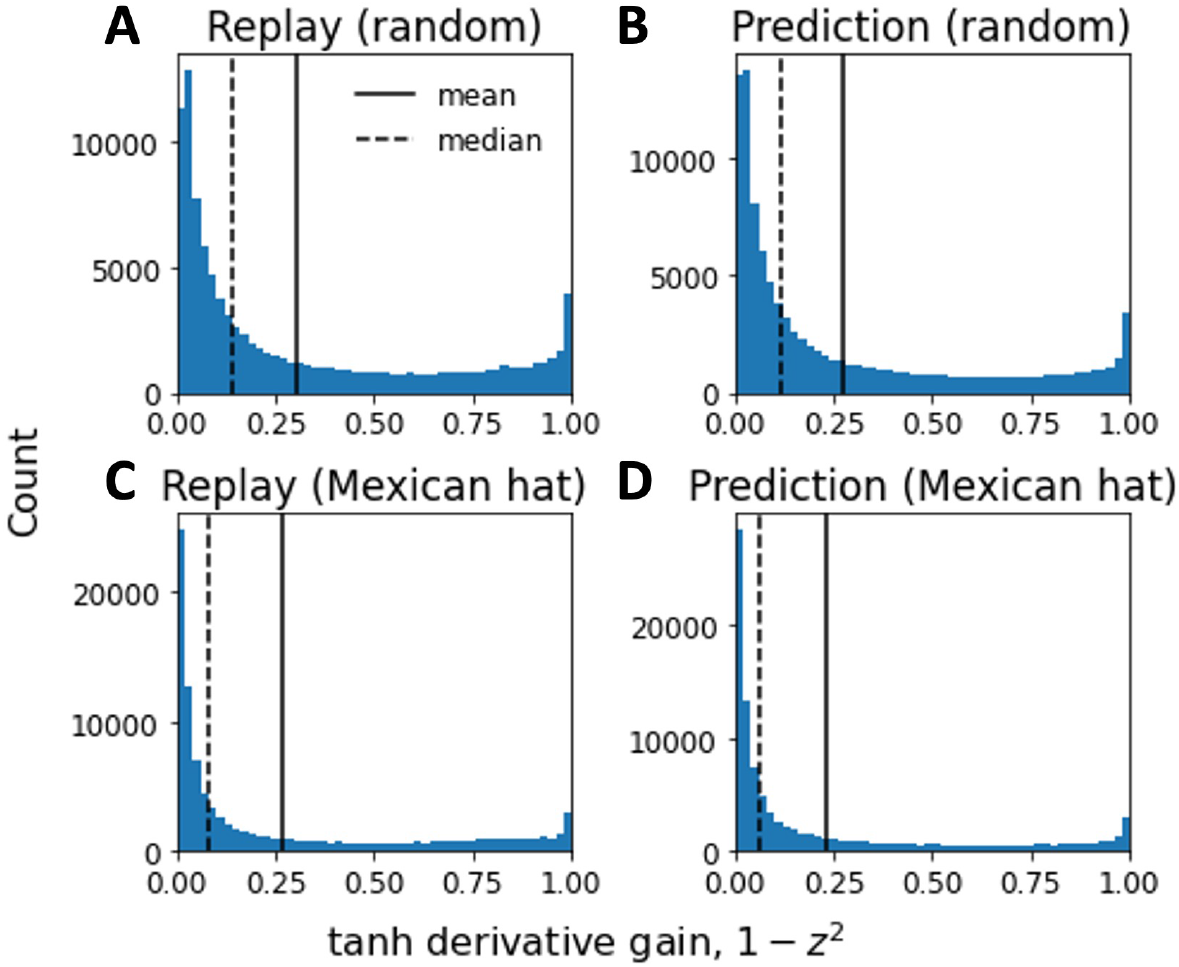
Effective gain of tanh nonlinearity during replay and prediction. Distribution of the instantaneous derivative 1 − *h*^2^ across hidden units and time evaluated for replay and prediction trajectories for networks with random and Mexican-hat initialization. Vertical lines indicate mean (solid) and median (dashed). The contraction factor (0.25-0.3) suggests effective spectral radius is reduced proportionally. In all conditions, the network operates in a moderately saturated regime, with mean gain of 0.2-0.3 and a substantial fraction of units exhibiting low gain (<0.1). Because the Jacobian is driven by 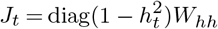, these results indicate that the effective dynamics correspond to a gain-scaled version of the recurrent matrix, contracting the spectrum relative to *W*_*hh*_.

At each time step, the network produces an *N*-dimensional population activity vector interpreted as a ring code. Outputs are rectified and normalized to form a population distribution,

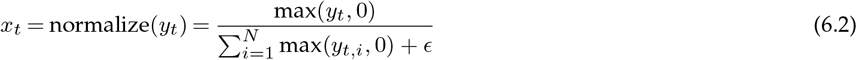

Each unit *i* is assigned a preferred angle *ϕ*_*i*_ = 2*πi/N*. Given the target ring distribution *p*_*t*_, the target angle is computed via its first circular moments

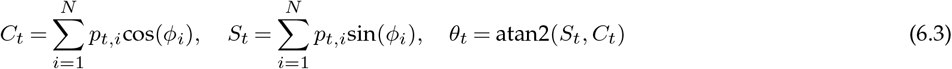

and represented as *Y*_*t*_ = [cos *θ*_*t*_, sin *θ*_*t*_]

Stacking all time points yields *X* ∈ ℝ^*T* ×*N*^ and *Y* ∈ ℝ^*T* ×2^. A ridge decoder is fit as

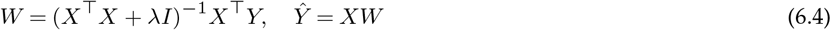

Performance is quantified by computing *R*^2^ separately for cosine and sine and averaging

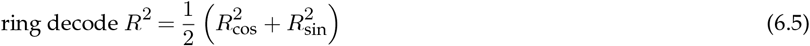

This metric measures time-resolved fidelity to the trained target phase *θ*_*t*_. It is not invariant to phase shifts, speed changes, or temporal offsets. For replay, it reflects spontaneous reproduction of the learned sequence under noise input. For prediction, it reflects accuracy in tracking the target trajectory given an initial drive. Values lie in (−∞, 1], with *R*^2^ = 1 indicating a correctly positioned, stable activity bump at all time steps.

#### (ii) Lag-1 autocorrelation

Lag-1 autocorrelation quantifies temporal dependence in the magnitude of model residuals. For each time step, we compute the residual magnitude

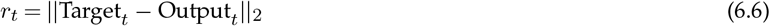

Then demeaning the resulting time series

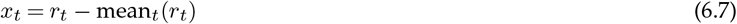

and compute the correlation between (*x*_1_, …, *x*_*T* −1_) and (*x*_2_, …, *x*_*T*_) (standard lag-1 autocorrelation).

Values lie in [−1, 1]. Zero indicates temporally uncorrelated (white-noise-like) residuals, positive values indicate smooth temporal persistence of errors, and negative values indicate alternating structure. High lag-1 autocorrelation reflects slowly varying residual magnitudes consistent with coherent underlying dynamics [103].

#### (iii) Mean resultant length (angular error concentration)

This metric quantifies how concentrated the circular angular error Δ*θ*(*t*) is over time between the true angle implied by the target ring distribution and the angle decoded from the model output [104]. It is also known as the mean resultant length, vector strength, or phase-locking value.

At each time step, target and model output activities are converted to angles, and the wrapped angular error Δ*θ*_*t*_ is computed.

Each error is represented as a unit vector

Let Δ*θ*_*b,t*_ be the wrapped circular error between the predicted and ground truth angle. We can define unit vectors as Then demean according to

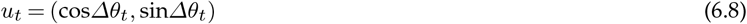

which are averaged over time

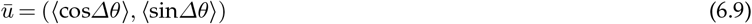

The mean resultant length is

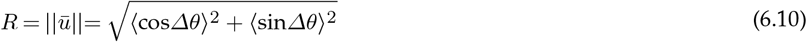

*R* ∈ [0, 1], where *R* = 1 indicates highly consistent angular errors (strong concentration near a single phase, typically zero) and *R* = 0 indicates uniformly distributed, random angular errors.

### (f) Symmetric STDP Analysis

The analytic form of the final weights is derived by convolving the Effective STDP Kernel (which includes the symmetry correction *ϵ*) with the Autocorrelation of the traveling Gaussian input. This is derived from the STDP learning kernel defined as

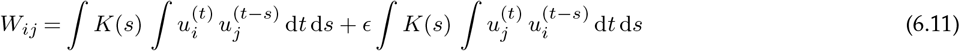

where

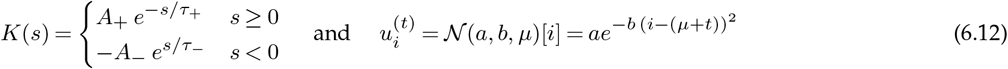

First computing the product 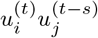

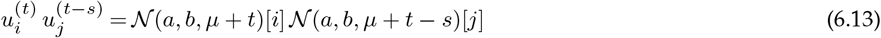

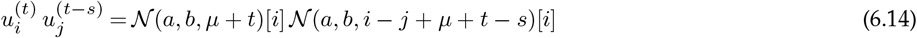

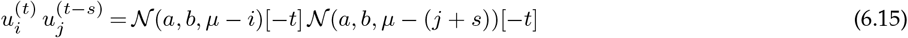

Using Lemma 1,

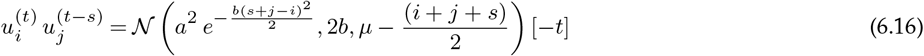

Now the autocorrelation,

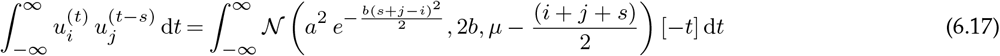

Using Lemma 2

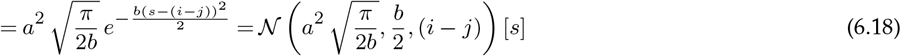

Computing 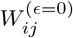 (the weight matrix without any symmetric correction) using autocorrelations

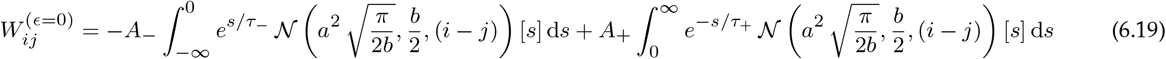

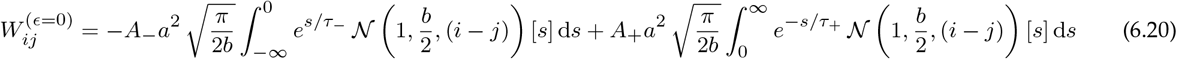

using Lemma 4

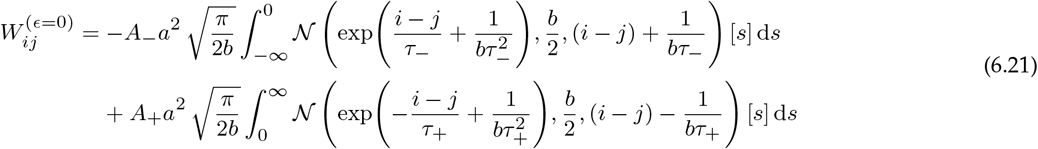

Let Δ = *i* − *j*,

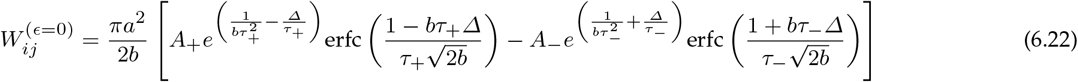

When the STDP has the symmetric correction (*ϵ*), we have:

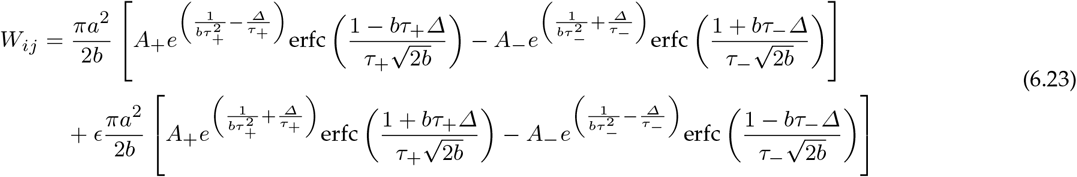

#### (i) STDP Eigenvalues

Eigenvalues control the dynamics the RNN has; however, they are difficult to compute directly from the equations for *W*_*ij*_. Instead, we compute it from the learning kernel in Equation **??** directly. Thus, the resultant *W*_*ij*_ is a Toeplitz matrix since *W*_*ij*_ is a function of Δ = *i* − *j*, which are asymptotically diagonalized by the DFT matrix. We have

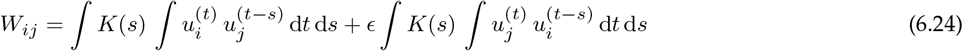

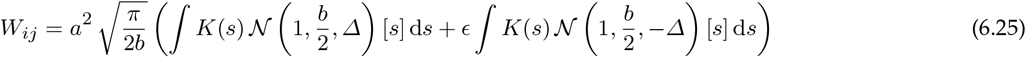

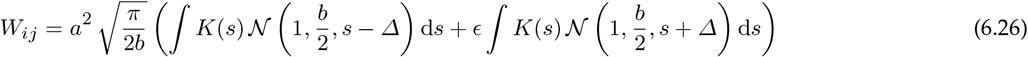

Using the convolution theorem, we have

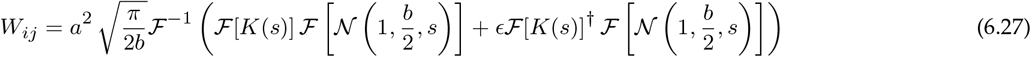

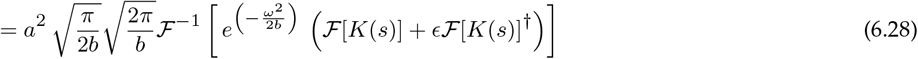

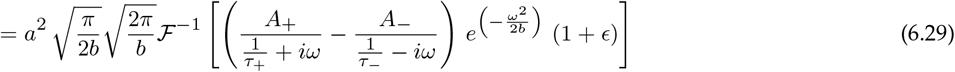

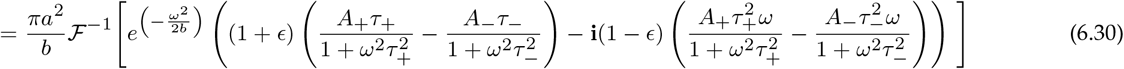

From this, we obtain the eigenvalue expression

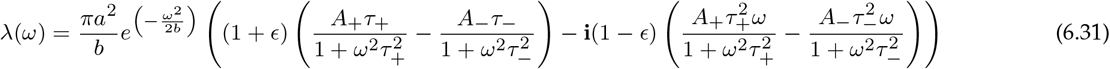

#### (ii) Useful STDP Lemmas

##### Lemma 1

(Product of Gaussian). *If* 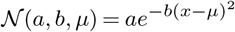, *then*

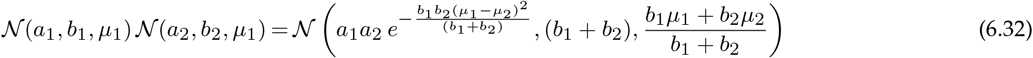

*Proof*. Let the product of the two Gaussian functions be defined as *f* (*x*) = *N* (*a*_1_, *b*_1_, *µ*_1_) *N* (*a*_2_, *b*_2_, *µ*_2_). By substituting the definition 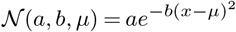, we obtain:

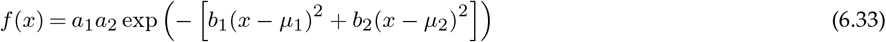

Let *Q*(*x*) denote the quadratic form in the exponent. Expanding the terms, we have:

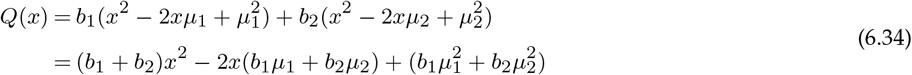

To express *Q*(*x*) in the standard form *b*_*c*_(*x* − *µ*_*c*_)^2^ + *K*, we define the combined precision *b*_*c*_ and combined mean *µ*_*c*_ as:

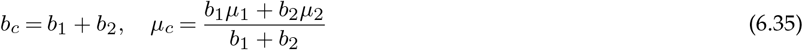

Rewriting the quadratic form by completing the square with respect to *x*:

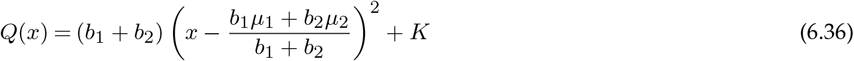

where the residual constant *K* is:

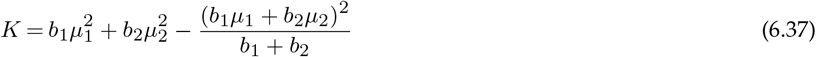

Simplifying *K* by finding a common denominator:

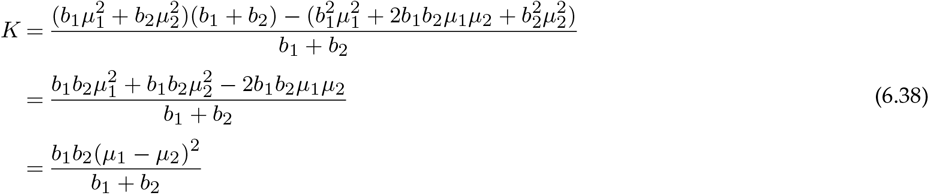

Substituting *Q*(*x*) = *b*_*c*_(*x* − *µ*_*c*_)^2^ + *K* back into the exponential function:

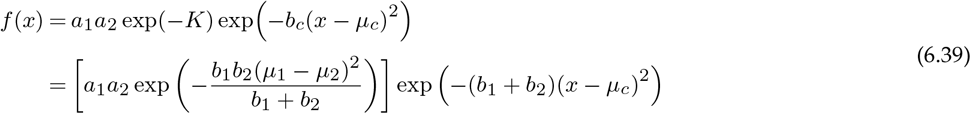

By inspection, this result corresponds to the function *N* (*a*_*c*_, *b*_*c*_, *µ*_*c*_).

##### Lemma 2

(Gaussian Integral). *If* 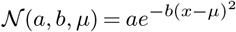, *then*

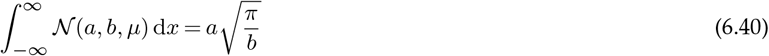

*Proof*. Let the integral be denoted by *I*. We aim to evaluate:

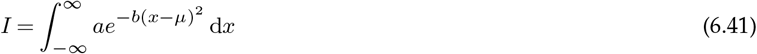

First, we factor out the constant amplitude *a*:

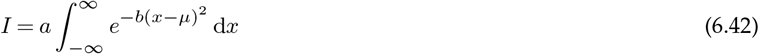

We perform a change of variables to simplify the exponent. Let 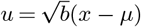. Consequently, the differential transforms as:

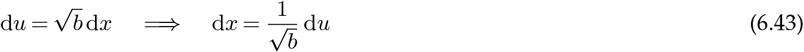

The limits of integration remain from −∞ to ∞. Substituting these into the integral:

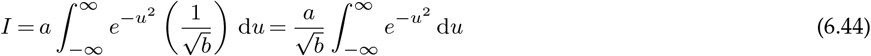

We recall the standard Gaussian integral identity:

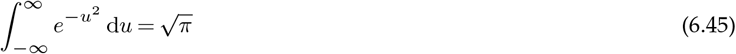

Substituting this standard result back into our expression for *I*:

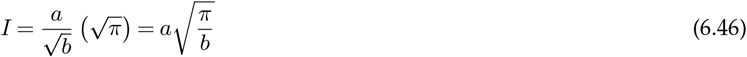

This completes the proof. □

##### Lemma 3

(Gaussian Integral Bounds). *If* 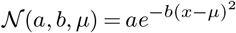, *then*

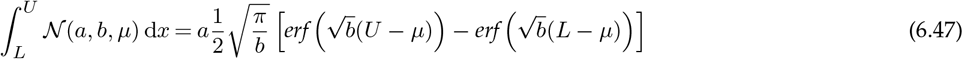

*Proof*. Let the integral be denoted by *I*. We aim to evaluate the definite integral:

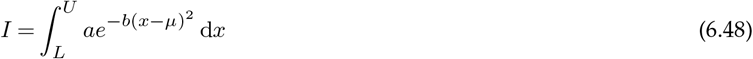

First, we factor out the constant amplitude *a*:

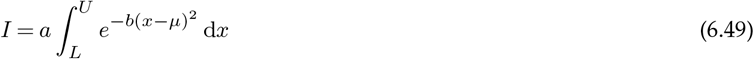

We perform a change of variables to standardize the exponent. Let 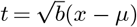. The differential transforms as 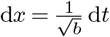. We must also transform the limits of integration:

- When 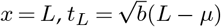.
- When 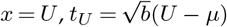.

Substituting these into the integral:

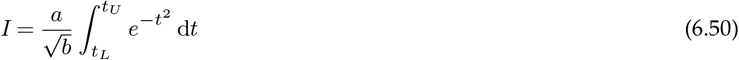

We recall the definition of the error function, erf(*z*), which is defined as:

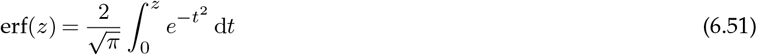

Using the property of definite integrals that 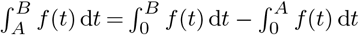, we can rewrite our integral in terms of the error function:

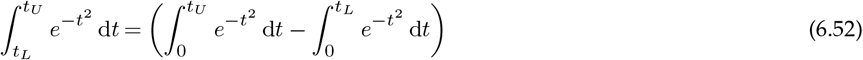

From the definition of erf(*z*), we have 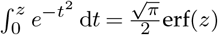. Therefore:

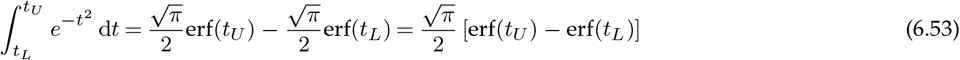

Substituting this result back into the expression for *I*:

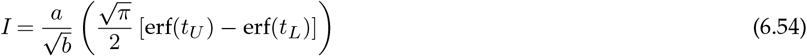

Rearranging terms and substituting the definitions of *t*_*U*_ and *t*_*L*_:

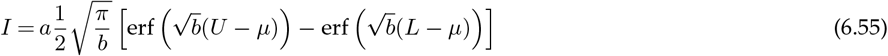

##### Lemma 4

(Gaussian Exp Product). *If* 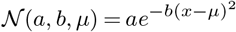, *then*

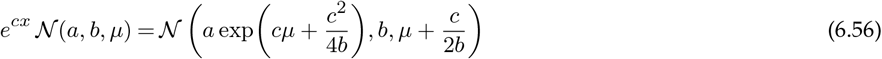

*Proof*. Let the product function be *f* (*x*) = *e*^*cx*^ *N* (*a, b, µ*). Substituting the definition of the Gaussian function:

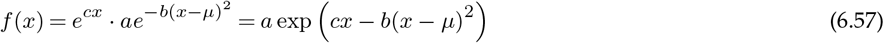

Let *E*(*x*) denote the exponent. We expand the quadratic term to group coefficients of *x*:

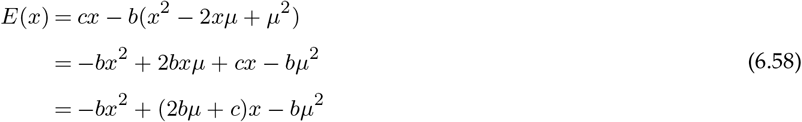

To restore the Gaussian form −*b*(*x* − *µ*_new_)^2^ + *K*, we factor out −*b* from the terms involving *x*:

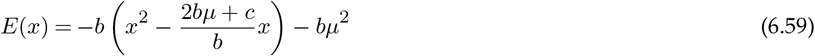

We define the new mean *µ*_new_ as half the coefficient of the linear term inside the parenthesis:

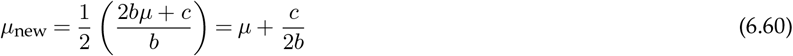

Completing the square inside the parenthesis by adding and subtracting 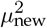:

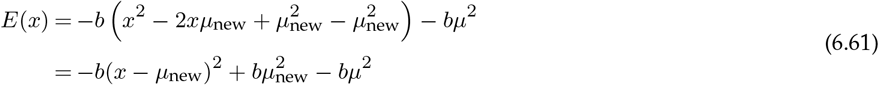

We simplify the constant terms (the residual) 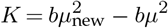:

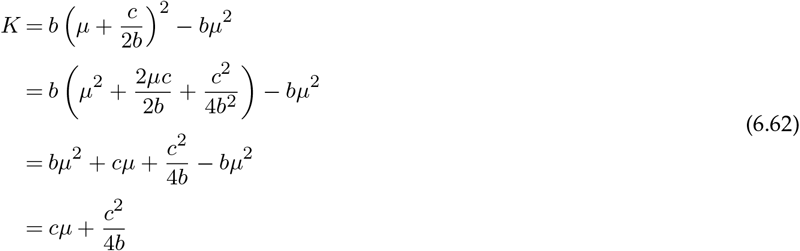

Substituting *E*(*x*) = −*b*(*x* − *µ*_new_)^2^ + *K* back into the original expression for *f* (*x*):

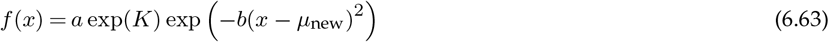

Substituting the values of *K* and *µ*_new_:

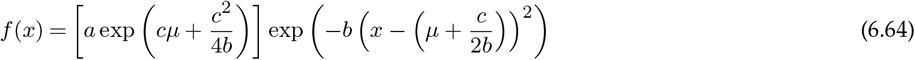

This matches the definition of *N*(*a*_new_, *b, µ*_new_). □

